# Brainwide organization of neuronal activity and convergent sensorimotor transformations in larval zebrafish

**DOI:** 10.1101/289413

**Authors:** Xiuye Chen, Yu Mu, Yu Hu, Aaron T. Kuan, Maxim Nikitchenko, Owen Randlett, Haim Sompolinsky, Florian Engert, Misha B. Ahrens

**Author notes:** equal contributions (Xiuye Chen, Yu Mu, Yu Hu, Aaron T. Kuan). senior authors (Haim Sompolinsky, Florian Engert, Misha B. Ahrens).

## Abstract

Simultaneous recordings of large populations of neurons in behaving animals allow detailed observation of high-dimensional, complex brain activity. However, experimental design and analysis approaches have not sufficiently evolved to fully realize the potential of these methods. We recorded whole-brain neuronal activity for larval zebrafish presented with a battery of visual stimuli while recording fictive motor output. These data were used to develop analysis methods including regression techniques that leverage trial-to-trial variations and unsupervised clustering techniques that organize neurons into functional groups. We used these methods to obtain brain-wide maps of concerted activity, which revealed both known and heretofore uncharacterized brain nuclei. We also identified neurons tuned to each stimulus type and motor output, and revealed nuclei in the anterior hindbrain that respond to multiple stimuli that elicit the same behavior. However, these convergent sensorimotor representations were only weakly correlated to instantaneous motor behavior, suggesting that they inform, but do not directly generate, behavioral output. These findings motivate a novel model of sensorimotor transformation spanning distinct behavioral contexts, within which these hindbrain convergence neurons likely constitute a key step.

## Introduction

Understanding the functional organization of the brain requires recording and interpreting activity from large populations of neurons. Recent advances in functional imaging technology, including the development of sensitive fluorescent reporters of neuronal activity (Akerboom et al., 2012; Chen et al., 2013) and fast imaging techniques (Bouchard et al., 2015; Fahrbach et al., 2013; Tomer et al., 2012), have made simultaneous recording across large brain areas possible in many animal models (Lemon et al., 2015; Peron et al., 2015; Prevedel et al., 2014; Schrödel et al., 2013). In the case of the larval zebrafish, the small size and transparency of the animal facilitates brain-wide optical interrogation of the nervous system (Ahrens et al., 2012; Friedrich et al., 2010; Orger, 2016; Orger et al., 2008; Portugues et al., 2014). Using light-sheet imaging techniques, the majority of the ∼100,000 neurons in the larval zebrafish brain can be imaged simultaneously at single-cell resolution in a behaving animal (Ahrens et al., 2013; Panier et al., 2013; Vladimirov et al., 2014).

Such large-scale simultaneous imaging has the potential to broadly reveal functional interactions between neurons across the brain and facilitate discovery of previously unobserved activity patterns. However, the scope of brain activity that can be understood is still limited to activity patterns actually explored by the brain during the experiment. To maximize this “neural space” explored by the brain, we extended the scope of whole-brain investigations by including a battery of different visual stimuli presented to each fish. Instead of focusing on only one behavior, we elicited phototaxis (Brockerhoff et al., 1995; Wolf et al., 2017), the optomotor response (Naumann et al., 2016; Orger and Baier, 2005), avoidance of visual looming stimuli (Dunn et al., 2016a; Lovett-Barron et al., 2017), and darkness responses (Burgess and Granato, 2007), as well as spontaneous behavior (Dunn et al., 2016b; Romano et al., 2015), in the same fish, while simultaneously monitoring fictive behavior (Ahrens et al., 2012; Masino and Fetcho, 2005). Eliciting multiple different behaviors leads to datasets rich in neural dynamics, facilitating the extraction of a wide variety of functionally coupled sets of neurons. This also uniquely enables us to examine the same neurons in various behaviorally relevant contexts, and obtain a more comprehensive view of their converging or diverging contributions across different sensorimotor pathways.

Inspecting and analyzing activity of ∼100,000 neurons poses a significant computational challenge, and requires the right tools. Many analysis approaches have been developed to interpret neural population activity data based on projections to low dimensional spaces (Cunningham and Yu, 2014; Freeman et al., 2014; Lopes-dos-Santos et al., 2013), regression (Feierstein et al., 2015; Miri et al., 2011a; Portugues et al., 2014; Wolf et al., 2017), noise correlation (Averbeck et al., 2006; Cohen and Kohn, 2011), and clustering (Romano et al., 2015, 2017). Here, we developed a combination of multiple analytical approaches to infer the organization of whole-brain activity during exposure to multiple stimuli, and quantified population dynamics both reflecting sensorimotor transformations and autonomous network activity across the brain.

First, we used regression analysis to quantify how closely the activity of each neuron is related to the stimulus inputs and motor output. However, in cases where visual stimuli robustly drive behavior, it may be hard to determine if cellular activity is more closely related to processing of stimuli or execution of motor commands. We therefore established an analysis for decomposing motor activity into stimulus-driven and stimulus-independent components by exploiting periodicities in the presented sensory stimulation and examining trial-to-trial variation in the motor output. In order to investigate functional relationships among neurons that are related to neither stimulus input nor motor output, we developed a density-based agglomerative clustering method for discovering groups of neurons with similar dynamics. Although cells were grouped solely based on functional activity, many of the resulting clusters were anatomically compact, revealing known as well as previously uncharacterized brain structures.

We also investigated how different sensory inputs diverge in the brain to represent different sensory features and then converge to trigger similar behavior (convergent stimuli) (Tononi et al 1998; Stein 1998; Hiramoto & Cline, 2009; Thill & Wilson 2016). We identified cells in the anterior hindbrain that are tuned to multiple different stimuli that induce right turns, along with their left turn counterparts. These cells are strongly correlated to the features of stimuli that drive a common behavior but are only weakly correlated to motor output, suggesting that they inform, but do not directly generate, behavioral output.

We implemented our analysis tools with a custom graphical user-interface (GUI) for interactive and flexible data exploration. The code, and all the data, will be made publicly available (after peer review of this manuscript), to enable community efforts for mining these rich datasets to discover deeper structure in whole-brain neuronal activity and its relation to visual stimuli and behavior.

## Results

### Whole-brain recordings of neuronal activity

Using a light-sheet imaging system reported previously (Vladimirov et al., 2014) (Fig. 1a), we recorded the activity of the majority of neurons in the larval zebrafish brain simultaneously, while presenting the battery of visual stimuli and recording fictive swimming behavior. For each fish, calcium imaging data was recorded at ∼2 volumes/sec for ∼50 min (2.11 ± 0.21 volumes/second; 6,800 ± 470 time frames, n = 18 fish) (Fig. 1b, Fig. S1e). The pan-neuronally expressed calcium indicator GCaMP6f (Chen et al., 2013) was localized to the cell nuclei by fusing it to the histone H2B protein (Vladimirov et al., 2014), which facilitates automatic cell segmentation performed via a template matching algorithm (Kawashima et al., 2016). We thus obtained the activity traces of ∼80,000 cellular ROIs per fish (8.0×10^4^ ± 1.6×10^4^ ROIs, n = 18; example traces in Fig. 1d), accounting for the vast majority of neurons in the brain with the exception of the most ventral part (Fig. S1a). During imaging sessions, fictive swim signals were detected by extracellular recordings of the descending axial motor neurons on each side of the tail (Ahrens et al., 2012; Masino and Fetcho, 2005). These recordings were reconstructed into fictive swim bouts, which were used to decode turns (Fig. 1d, bottom bar; **Methods**) (Ahrens et al., 2013; Dunn et al., 2016b).

**Figure 1.**
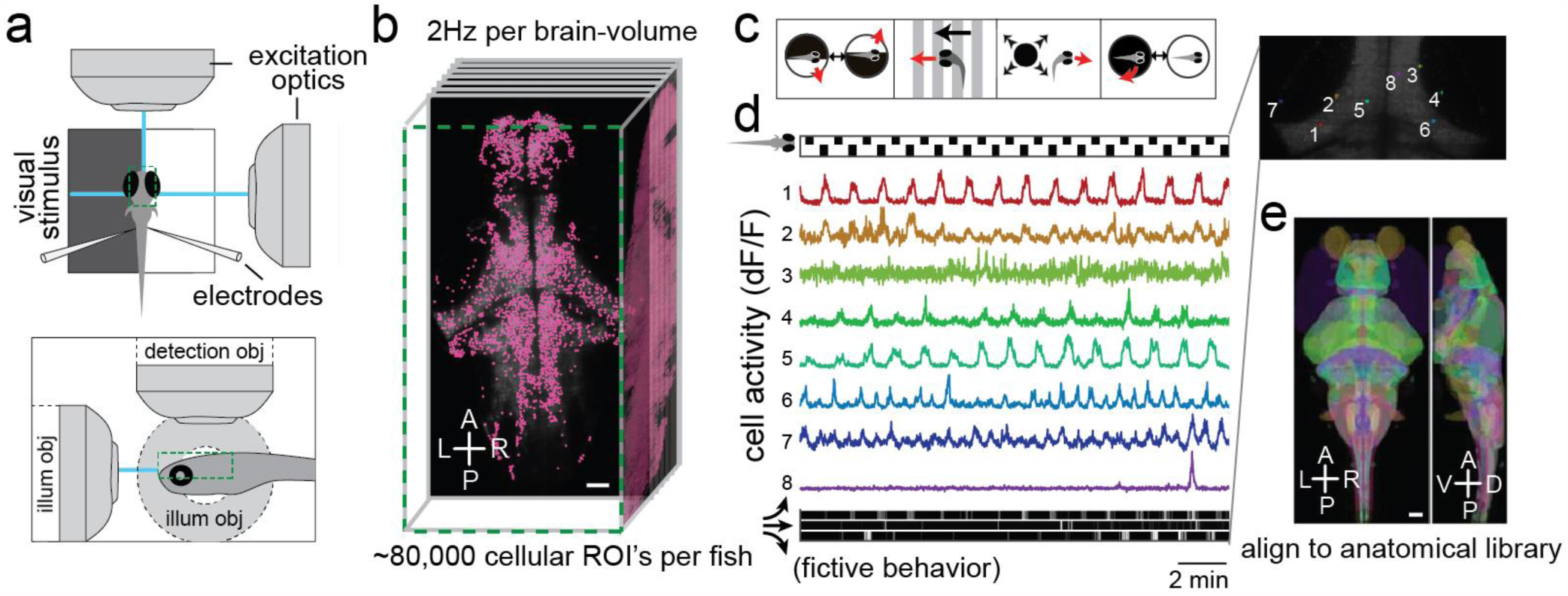
Whole-brain recordings of neuronal activity. (**a**) Schematic of experimental setup for fictive swimming combined with light-sheet imaging (see Methods). (**b**) Illustration of functional dataset format. Whole-brain volumes were imaged at ∼2 volumes/sec for ∼50 min (2.11 ± 0.21 volumes/second; 6,800 ± 470 time frames, n = 18 fish). The activity traces of individual neurons were automatically extracted for 8.0×10^4^ ± 1.6×10^4^ cellular ROI’s per animal (n = 18). Scale bar, 50 µm. (**c**) Illustration of visual stimuli presented during functional imaging. Four stimulus paradigms from left to right: phototactic stimulus (phT), moving stripes (Optomotor response or OMR), expanding dot (looming or visual escape response), and dark flashes. (**d**) Example neuronal activity of single neuron ROI’s within an imaging plane in the tectal region in the midbrain. The stimulus (phototactic stimuli alternating with white background) is illustrated with a bar above the calcium traces, and the recorded fictive behavior is plotted at the bottom of the panel, with three plots indicating left turns, forward swims and right turns from top to bottom. (**e**) Image stacks were registered to the Z-brain reference brain atlas (Randlett et al., 2015) containing ∼300 labels of anatomical regions. Scale bar, 50 µm.

Blocks of visual stimulus patterns associated with the following behavioral paradigms were projected onto a screen below the fish during imaging (Fig. 1c). (1) Phototaxis. Larval zebrafish are attracted by light and are averse to darkness, and use spatial differences in luminance to guide their navigation (Brockerhoff et al., 1995; Wolf et al., 2017). We presented a half-field dark stimulus on either the left or the right side, separated in time by a whole-field white baseline. (2) The optomotor response. This is a position-stabilizing reflex to whole-field visual motion, in which fish turn and swim in the direction of perceived motion (Naumann et al., 2016; Orger and Baier, 2005). The stimuli used were whole-field stripes moving in different directions. (3) The visual escape (looming) response. In free-swimming fish, expanding discs elicit an avoidance response; although in tethered preparations fish often exhibit freezing behavior, there is still a strong sensory-related response in large areas of the brain (Dunn et al., 2016a; Lovett-Barron et al., 2017). The looming stimuli were presented from either the left or right side of the fish. (4) The dark-flash response. This reflex is characterized by large-angle turns in response to sudden darkening of the environment (Burgess and Granato, 2007; Chen and Engert, 2014). Alternating whole-field dark and bright stimuli were used for this stimulus block. (5) Spontaneous behavior. In the absence of visual stimuli, fish spontaneously swim in alternating sequences of repeated turns to the left and to the right (Dunn, Mu et al. 2016). For this block, fish were imaged under homogeneous background illumination.

In order to make anatomical comparisons across experiments with different fish, we registered the functional imaging stacks to each other (Fig. S1b,c), and also to the Z-Brain atlas for larval zebrafish (Randlett et al., 2015) that contains molecular labels and definitions for known anatomical regions (Fig. 1e, Fig. S1d). To enable intuitive data analysis, we developed a custom interactive platform in MATLAB (Fig. S1f, code and data available online) to efficiently explore this data in both functional and anatomical contexts. All of the analysis presented here was performed using this GUI platform and associated scripts.

### Identification of sensory and motor related neurons

A first step towards understanding the brain’s sensorimotor transformations is identifying cells whose neural activity closely tracks the stimulus input or motor output. This type of analysis provides a survey of the cells that are likely involved in a given sensorimotor transformation. With whole-brain data including a broad range of stimulus types, one can simultaneously obtain comprehensive tuning maps for multiple stimulus types and motor outputs, which allows for comparing neural pathways for different stimuli. We describe below a highlight of results from the individual stimulus or motor maps and from comparing these maps. We first constructed left and right motor tuning maps by regressing whole-brain activity against fictive recordings of motor behaviors, from which turn direction can be reliably decoded (Dunn et al., 2016b)(Fig. 2a,b,c, **Methods**; see also Fig. S2g for left/right/forward maps). Despite the simplicity of the regression analysis (and limitations that we will discuss in the next section), these maps show a dense, lateralized and highly concerted neural population in the hindbrain (Fig. 2c), which is consistent with results from previous studies during spontaneous fictive swimming (Dunn et al., 2016b). A combined motor map incorporating results for multiple animals reveals well-known motor-related anatomical landmarks with remarkable precision, including the RoV3, MiV1 and MiV2 neurons (Fig. 2c, upper right inset) (Orger et al., 2008; Randlett et al., 2015).

**Figure 2.**
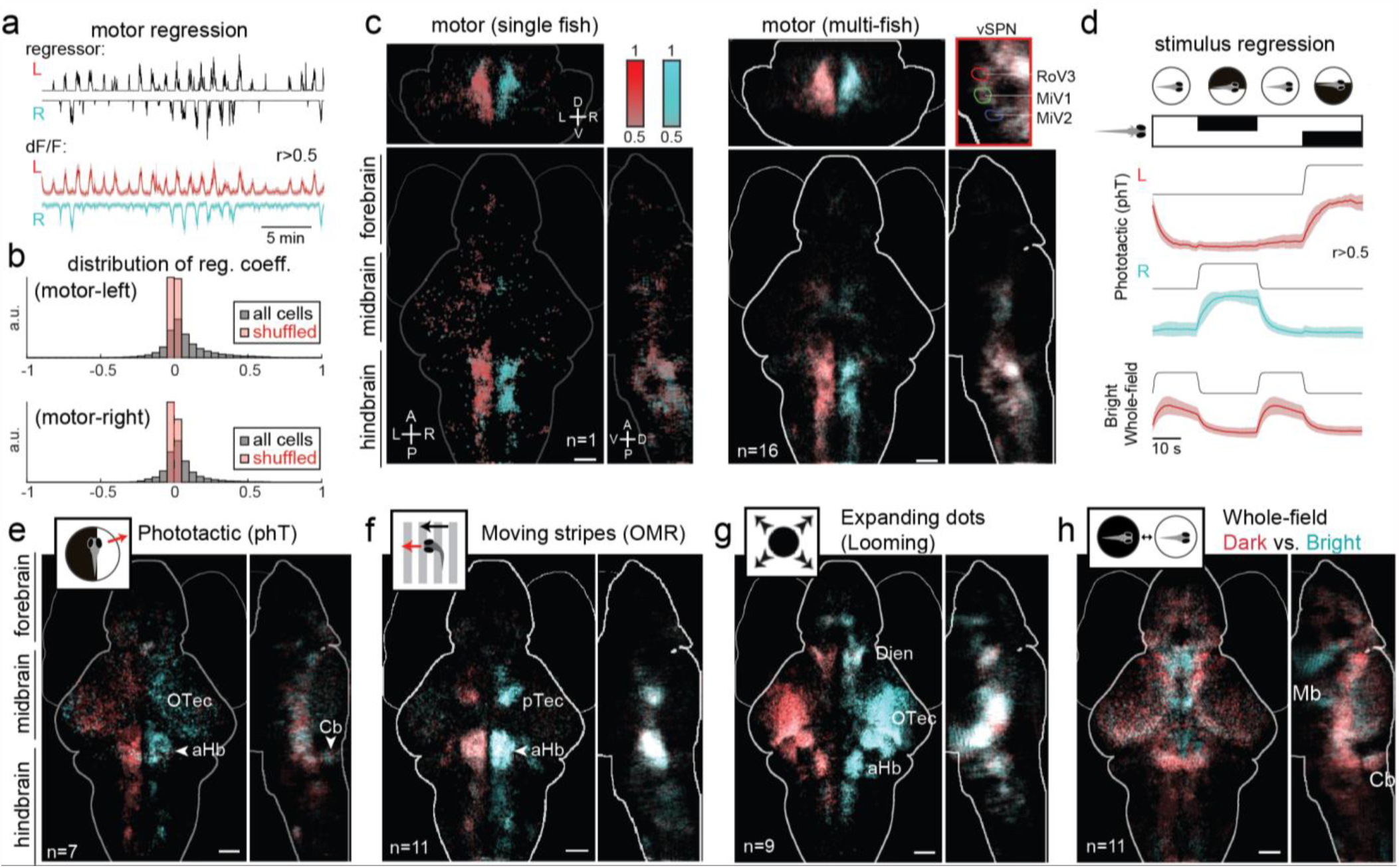
Identification of sensory and motor related neurons. (**a**) Motor regression: motor regressors (for left, right turns, respectively) were constructed by convolving the processed fictive swimming traces with an impulse kernel. Brain activity from all cellular ROI’s is regressed against these regressors. The dF/F traces show the mean±SD for all ROI’s with r>0.5 for a single fish. (**b**) Distribution of regression coefficients for all cells in the example fish, as compared to data with shuffled time traces. **(c)** Left: anatomical map of all ROI’s with r>0.5 to either leftwards (red) or rightwards (cyan) motor regressors, from the same fish. Right: Average response maps across fish (n=16). Inset: activity in the proximity of the hindbrain spinal projection neurons that control turning (Orger et al., 2008), namely RoV3, MiV1 and MiV2 neurons; masks from Z-Brain Atlas. (**d**) Stimulus regression, using phototactic stimulus as example. Fish were shown a periodic stimulus during imaging that consists of leftwards and rightwards phototactic stimuli separated by a whole-field bright background. The stimulus regressors (black) are constructed by convolving a binary step function with an impulse kernel of GCaMP6. The colored traces show the dF/F (mean±SD for all ROI’s with r>0.5) for the same example fish as in (a). (**e, f, g, h**) Average stimulus response maps for (**e**) phototactic stimuli (phT), (**f**) moving stripes (Optomotor response, or OMR), (**g**) expanding dots (looming or visual escape response), and (**h**) whole-field dark versus bright (dark-flash response). Scale bars, 50 µm.

Next, we constructed sensory regressors (Fig. 2d, see Methods) and maps for each of the specific types of stimuli in this experiment, including phototactic (phT, Fig. 2e), optomotor (OMR, Fig. 2f), looming (Fig. 2g), and dark-flash (Fig. 2h) stimuli. We first describe the results from the phototactic tuning maps, which show (left/right) symmetric/lateralized and anatomically dispersed activity in the midbrain optic tectum, as well as active populations in the cerebellum (Fig. 2e). The dominant responses to phT in the optic tectum come from the contralateral eye, which indicates that they respond to the dark half of the stimulus. These OFF responses in the optic tectum are also ipsilateral to the activity in the hindbrain, which is more motor-related (Fig. 2c). In accordance with unilateral laser ablation experiments (Burgess et al., 2010), this suggests that the OFF pathway is also responsible for the lateralized anterior hindbrain activity. Indeed, regression maps based on individual components of the phototactic stimulus (bright whole-field, bright half-field, and dark half-field/phototactic, Fig. S2h,i,j,k and Fig. 2e) further suggest that phototaxis behavior is more strongly driven by the dark half-field than the bright half-field component, because only the dark half-field/phototactic map contains neurons in the more caudal hindbrain that overlap with the motor tuning map.

The moving stripes stimulus elicits strong and robust responses in the pretectum and anterior hindbrain, again consistent with previously reported data (Fig. 2f, see also Fig. S2d for OMR forward vs backward) (Naumann et al., 2016). The looming stimulus evokes striking and bilaterally symmetric activity patterns in large areas of the hypothalamus, in stereotypical locations within the forebrain and diencephalon, as well as in the hindbrain (Fig. 2g). A comparison between the OMR and looming activity maps (Fig. 2f,g and Fig. S2l) shows that distinct areas of the anterior hindbrain are activated. In comparison, there is significantly more overlap between the maps for OMR and phototaxis (Fig. 2e,f, see also Fig. S2a). This suggests that sensory–motor processing pathways for OMR are more similar to phototaxis than escape responses. One caveat, however, is that while the OMR paradigm works robustly in tethered preparations, animals respond to looming stimuli much less frequently (see Fig. S2e,f for motor maps during OMR vs. looming), which may partially account for the distinct activity maps. The activity maps for dark and bright whole-field stimuli (Fig. 2h) are also bilaterally symmetrical and quite distinct from the other maps. Interestingly, most of the activation unique to the whole-field stimuli (i.e., not seen in the other 3 maps) is related to the bright stimulus and is located close to the midline in the midbrain area. This points to a specific region-of-interest for further analysis on the circuits involved in processing whole-field bright flashes. In summary, as a first-pass analysis in identifying neurons involved in sensorimotor transformations, the whole-brain stimulus and motor regression maps can already offer insights for distinguishing models and forming hypothesis for detailed analysis.

### Dissection of sensorimotor components based on trial-to-trial variations

If the behavioral output were entirely independent of the stimulus input, then the separate stimulus/motor regression analyses above would provide an adequate description. However, for stimulus-evoked behaviors, there is generally a strong correlation between the stimulus and motor regressors. A simple regression analysis can therefore mix up stimulus and motor-related neuronal activity, limiting its effectiveness beyond identifying neurons with very high regression coefficients.

To address this issue, we took advantage of the fact that the stimuli presented in our dataset are exactly periodic. Specifically, we expect the stimulus-driven component of motor activity to be approximately periodic (following the stimulus), and other independent motor activity to be spontaneous and aperiodic. The activity of each neuron can thus be decomposed into trial average (periodic) and residual (aperiodic) components, representing stimulus-driven and stimulus independent activity (Fig. 3a, columns). The trial average activity can be thought of as a generalization of stimulus regressors. Indeed, clusters of cells can be found with many types of periodic activity patterns (Fig. 3b), including but not limited to the simple regressors used previously in Fig. 2. By selecting cells based on the amplitude of the trial average, one can map generally the most stimulus-responsive cells (as opposed to those exhibiting a particular stimulus pattern, as in Fig. 2) (Fig. 3c, see Fig. S3c for least stimulus-responsive cells). In this generalized sensory map, we find more cells broadly distributed in the optic tectum, forebrain, anterior hindbrain and cerebellum.

**Figure 3.**
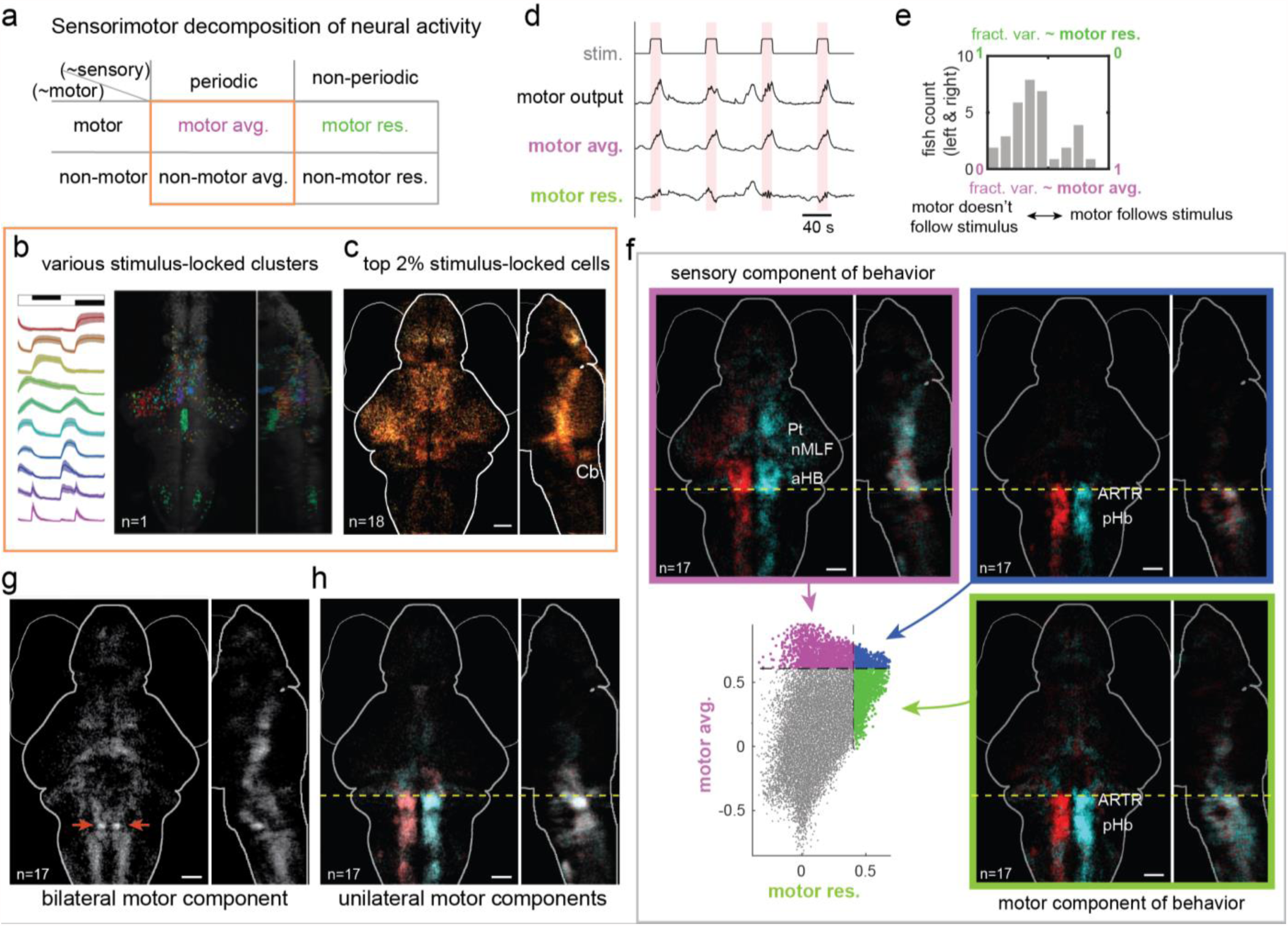
Dissection of sensory-motor components based on trial-to-trial variations. (**a**) Schematic illustration of sensory-motor decomposition of neuronal activity into periodic/aperiodic and motor/non-motor components. (**b**) Various functional clusters of cells with large periodic component for phototactic stimuli (cells selected for low variance across repetitions, curated k-means clusters from a single fish). Left: dF/F (mean±SD). (**c**) Average map of cells with most highly periodic activity. Cells are ranked by the variance explained by their periodic component. The top 2% of cells for each fish are selected, and the average from n=18 fish is shown. (**d**) Example decomposition of the motor output into periodic (motor avg.) and residual (motor res.) components. (**e**) Histogram of the fraction of variance explained by the periodic component of the motor output (n=17 fish, left and right motor outputs calculated separately). (**f**) Dissection of brain-wide trial-average and residual motor activity. Lower left panel: Scatter plot of regression coefficients for all cells with respect to motor avg. (y axis) and motor res. (x axis) regressors, for an example fish in relation to the left-side motor output. Top left, purple box: map showing cells ranking in the top 2% for motor avg. regression only. Bottom right, green box: map showing cells ranking in the top 2% for motor res. regression only. Top right, blue box: map showing cells ranking in the top 2% for both motor avg. and motor res. regressions. (**g-h**) Bilateral versus unilateral motor components. Analogously to subtracting the trial average from the full activity traces, here we obtain the frame-by-frame average of the left and right motor outputs (bilateral component) and calculate left-right residuals (unilateral components) for the left and right side, respectively (see Fig. S3e). (**g**) Cells ranking in the top 2% for bilateral regression only. Note the bilateral and dense cluster of cells at the red arrow locations in Rhombomere 5. (**h**) Cells ranking in the top 2% for unilateral regressions only. Dotted yellow lines: boundary between Rhombomeres 2 and 3. Scale bars, 50 µm.

Similarly, the motor outputs can also be decomposed into trial average and residual components, which represent stimulus-driven motor activity (motor avg.) and independent motor activity (motor res.) (Fig. 3d,e). To faithfully represent the relevant neural dynamics, we defined the motor outputs (left and right) as the activity of selected neurons that are most highly correlated to the fictive recordings (**Methods**; Fig. S3a,b). To explore the distinction between the two components of motor output, we plotted the regression coefficient of each cell to motor avg. and motor res. regressors in a two-dimensional space (Fig. 3f, lower left, regressions to left-side motor output for an example fish). We then plotted the anatomical locations of neurons that ranked highly (top 2%) in motor avg. only, motor res. only, or both (Fig 3f, purple, green, and blue boxes, respectively). In this analysis, we found that constructing maps based on cell ranking, rather that absolute value of motor residual components, produced maps that are more robust when comparing across animals. The motor res. map (Fig. 3f, green box) is similar to the motor regression map shown in Fig. 2a, but contains neurons more exclusively in the hindbrain, consistent with the idea that selecting for the motor residual filters out motor activity that is driven by the stimulus. The motor avg. map (Fig. 3f, purple box) shows significant populations of neurons in the midbrain, which are highly consistent with anatomical regions (pretectum and nMLF) that are known to be central for the processing of these visual stimuli. The map also includes cells in the anterior hindbrain region (aHB), consistent with previous observations for OMR stimuli (Naumann et al., 2016). The intersection map (Fig. 3f, blue box) includes cells almost exclusively posterior to the rhombomere 2/3 boundary in the hindbrain. Compared to the motor res. map, cells here are more tightly clustered in the “hindbrain oscillator” (Ahrens et al., 2013; Wolf et al., 2017), also known as the “anterior rhombencephalic tuning region (ARTR)” (Dunn et al., 2016b), suggesting that this brain area may play an important role in the sensorimotor transformation (see also Fig. S5a,e).

Our approach of dissecting motor activity into trial average and residual components can also be analogously applied along the laterization dimension between left and right motor outputs (Fig. S3e). Specifically, we extracted the average response across left and rightward stimulus presentations to form a bilateral component. Subtracting this bilateral component from the left and right motor outputs yields unilateral residual components. These unilateral motor outputs are more sensitively tuned to activity associated with turns in a particular direction, as opposed to forward swimming. Similar to the previous analysis, we produced tuning maps for cells ranking highly (top 2%) either in the bilateral (average) (Fig. 3g) component only, or the unilateral (residual) components only (Fig. 3h, see also Fig. S3f). These maps uncover more anatomical subtleties: the bilateral map features a tight pair of clusters in Rhombomere 5 near the MiD2 spinal projection neurons that to our knowledge has not been previously characterized, and the unilateral maps reveal a prominent pattern of contralateral correlations across the Rhombomere 2/3 boundary.

### Whole-brain functional clustering

Although the sensorimotor system is central for understanding the functional organization of the brain, many neurons in the brain exhibit coherent activity that is not strongly correlated with either the visual stimuli or motor outputs. This should not be surprising, particularly because (1) we did not exhaust the sensory or behavioral repertoire of the animal, and our recorded motor correlates are restricted to swimming; (2) brain activity may be modulated by internal dynamics and brain states not directly related to stimuli or actions (Flavell et al., 2013; Kawashima et al., 2016); and (3) the larval zebrafish brain is in a stage of rapid development in which young neurons are continuously incorporated into existing circuits (Boulanger-Weill et al., 2017). Thus, to generate a more comprehensive map of the functional organization of the larval zebrafish brain, we clustered neurons based on the correlations between their activities. This greatly reduces the dimensionality of the data without sacrificing biological interpretability (e.g. compare with spatial ICA in Fig. S4-1l-m).

We devised an unsupervised density-based agglomerative clustering algorithm that groups neurons into functional clusters based on the functional activity alone (regardless of their anatomical location, see Methods and Fig. S4-1a; example clusters in Fig. 4a,b for 1 example animal). The algorithm has a tunable threshold regulating how correlated the activity of cells within the same cluster should be (**Methods**). Cells that are not correlated to any cluster above this threshold remain unclustered. The value of the threshold affects the clustering as follows: as the threshold is increased, the activity of cells within clusters becomes more similar, and the clusters become more robust to a cross-validation test (Fig. S4-1d,e), but fewer cells are included in clusters, and the number of clusters is reduced. As the threshold is decreased, more cells are included in clusters, but some anatomically distinct clusters merge together into larger, more loosely associated groups of cells. For the ensuing analysis, we have empirically chosen a threshold (0.7) that balances this tradeoff. With this criterion, each fish contains ∼100-150 functional clusters, and each major brain area contains at least a few clusters (Fig. 4b). We find that the number of cells included in each cluster spans three orders of magnitude (Fig. 4d). Note that this size heterogeneity is not an artifact of poorly defined clusters, as the activity within each cluster is highly concerted (Fig. 4e; examples in Fig. 4a).

**Figure 4.**
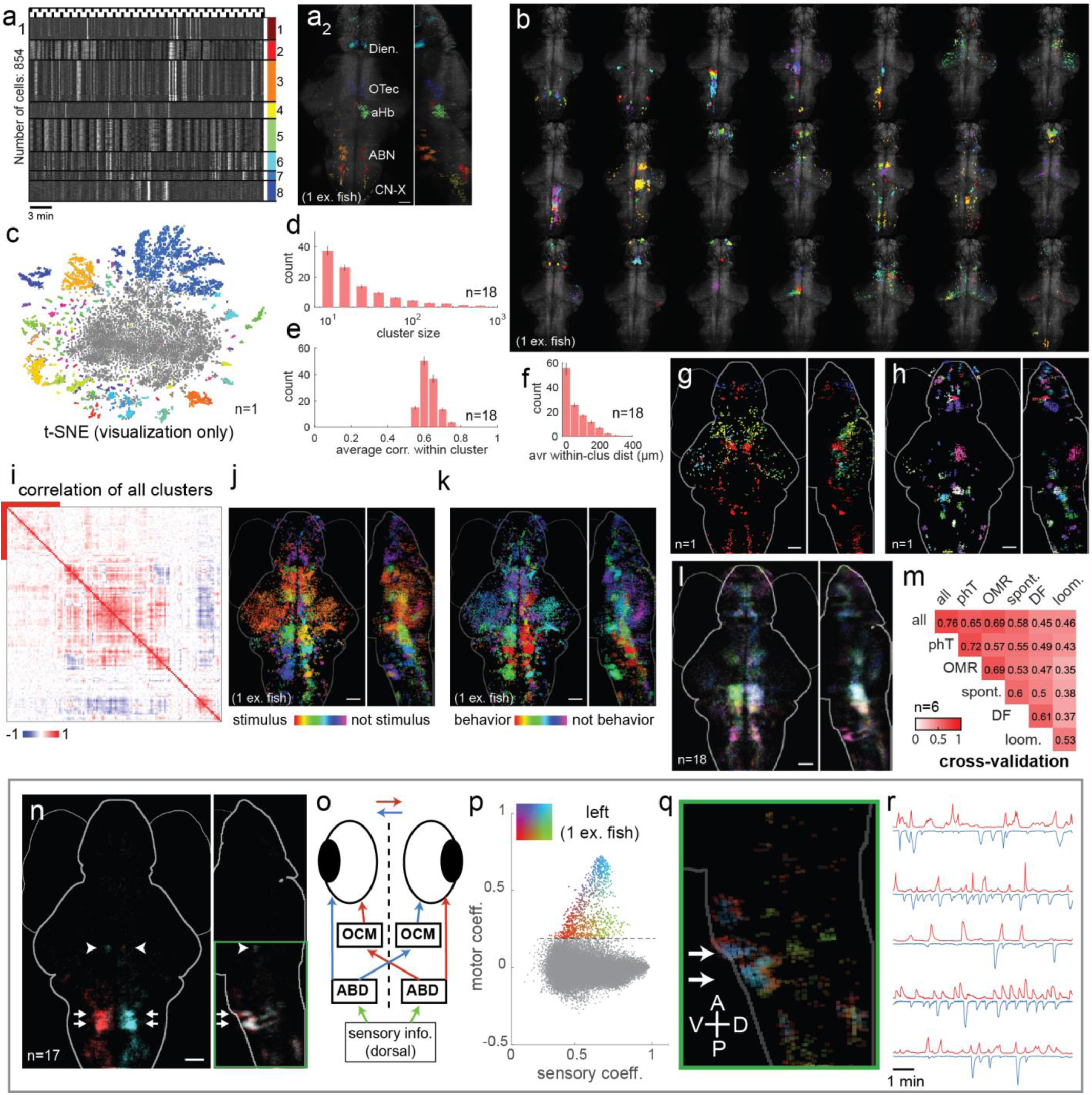
Whole-brain functional clustering. (**a**) A diverse collection of automatically identified functional clusters from one example fish. After whole-brain clustering, the most dissimilar clusters were selected based on their hierarchical ranking. Each color represents one cluster. a1: functional activity profiles: normalized ΔF/F for each cell is plotted along the horizontal axis. a2: corresponding anatomical map. (**b**) Full set of individual clusters as z-projections (from the same fish), in groups of 6 per projection (for distinguishable color assignments). (**c**) Visualization of all clusters within one fish with t-SNE. For color assignment, the total of 139 clusters (6,499 cells) were ordered by hierarchical clustering, and adjacent shades of hsv colors are assigned based on the resulting leaf order. Gray points represent cells that did not pass the clustering criterion i.e. not assigned to any cluster (1:10 down-sampled for clarity). (**d**) Histogram of size distribution of all clusters, pooled across fish (mean±SEM). (**e**) Histogram of average correlation between cells within clusters, pooled across fish (mean±SEM). (**f**) Histogram of average within-cluster anatomical distance, pooled across fish (mean±SEM). (**g**) Examples of anatomically dispersed clusters that are difficult to identify based on anatomical location, as they are intermixed with other clusters. (**h**) Examples of clusters that anatomically isolated. (**i**) Correlation matrix of all clusters from a single fish, ordered so that most similar clusters are adjacent. Submatrix indicated by red bar are putative olfactory bulb neurons (see Fig. S4-2f). (**j**) Clusters for an example fish ranked from stimulus-related (periodic, as in Fig. 3c, red) to not stimulus locked (aperiodic, purple). (**k**) Clusters for an example fish ranked from motor-related (high regression coefficient to motor res., as in Fig. 3f, red) to not motor-related (low regression coefficient, purple). (**l**) Average map of clusters conserved in anatomical space. Each of these clusters are selected for having anatomically corresponding clusters in at least 6 other fish (out of the 18 fish assessed, see Methods). Clusters are ranked and colored within fish as in (k). (**m**) Two-fold cross-validation (as in Fig. S4-1e) between multiple sets of stimuli within each fish (n = 6 fish), with scores indicating the fraction of cells in matched clusters over the total number of cells. Each fish in the analysis has been presented with all 5 different stimuli. The “all” category uses the combined data from all 5 stimulus periods. Color coding is the same as number labels. Scale bars, 50 µm. (**n**) Putative abducens nucleus (ABD) network for the control of eye-movements. Arrows: anterior and posterior clusters of the ABD map to Rhombomeres 5 and 6, respectively. Arrowheads: oculomotor nucleus (OCM) clusters. (**o**) Illustration of the proposed eye-control circuit. Red/blue indicates control of rightward/leftward eye movement. (p-q) Two-dimensional sensory-motor mapping as in Figure 3f, except using the average response of the abducens nucleus (curated ROI) as eye-movement regressors (instead of the tail-movement motor output). (**p**) Analysis of the sensory and motor activity of one example fish. Horizontal axis: stimulus component (square root of variance explained by periodic component of activity); vertical axis: motor component (regression coefficient with left motor res., as in Fig. 3d); Top 1000 motor ranked cells are plotted in colors, corresponding to (q). (**q**) Anatomical map of cells shown in (p). See Fig. S4-2g for corresponding figures for right motor res. (**r**) Putative eye-movement traces, extracted from averaging the neural activity of highly concerted functional clusters representing the ABD nucleus. Red/blue: left/right.

We also examined the anatomical structure of these functionally identified clusters. Since the clustering uses no anatomical knowledge as input, emergent anatomical patterns in the clusters can reveal underlying organizational relationships between structure and function within the brain. Interestingly, most clusters are anatomically compact (Fig. 4f and Fig. S4-1c), suggesting that functionally related neurons are often organized into small brain nuclei (Fig. 4h). However, the spatial extent of clusters varies significantly (Fig. 4f tail), including prominent examples of spatially dispersed clusters that are spatially intermingled with other clusters (Fig. 4g).

Many of the clusters exhibit substantial correlations with other clusters (Fig. 4i, correlation matrix across clusters). To capture the complex relationship between clusters, we also performed hierarchical ordering of the cluster centers (Fig. S4-1b,f). The blocks or branches of this hierarchy reveals additional functional organization on multiple broader scales (Fig. 4i, red block corresponds to the putative olfactory bulb neurons, see Fig. S4-2f). Similar to the analyses performed in Figs. 2 and 3, we situated clusters in sensory and motor dimensions by ranking them based on stimulus or behavior-related components (Fig. 4j,k). Regions that are most tuned to stimulus and motor are consistent with the cellular activity maps shown previously (Fig. 2c and Fig. 3f). However, these cluster-based maps also highlight clusters (for example in the forebrain) that lie on the lower end of both the stimulus and motor ranking maps, demonstrating coherent activity patterns that are not tied to either stimulus or motor.

One might expect that many clusters would appear in similar anatomical locations across different individual animals. However, it is also possible that some functional clusters appear at different locations in different animals. To investigate this, we screened for clusters that have a similar anatomical counterpart in several individual fish. About half of all clusters are anatomically conserved according to our criteria (Fig. S4-1h; see **Methods** and Fig. S4-1g for criteria). A population-averaged map of these anatomically conserved clusters (Fig. 4l) highlights the anterior hindbrain and ventral forebrain, among other regions, while clusters such as those in the optic tectum are absent, consistent with previous observations that stereotypy in the tectum is low (Portugues et al., 2014; Randlett et al., 2015).

Unsupervised clustering faces the challenge of the lack of ground truth for validating the derived clustering structure. To address this, we first visualized our clustering result independently using an unsupervised data visualization method, t-SNE (i.e. the location of the neurons in the t-SNE plot are not informed by the clustering results) (Van Der Maaten and Hinton, 2008). The functional clusters are represented as isolated islands in the periphery of the t-SNE plot (colored dots), while unclustered cells (i.e., with noisy or idiosyncratic activity below the density threshold) congregate in the center (grey dots) (Fig. 4c). This suggests that our clustering results and the choice of clustering threshold are generally appropriate.

To more systematically evaluate the clustering results, we performed a cross-validation analysis that compares clusters obtained using one half of the time-series data versus the other half. We find that 76% of clusters are reliably obtained from both halves of the time-series data (Fig. 4m, top left; **Methods** and Fig. S4-1e,n). A similar cross-validation method can be applied to investigate how cluster relationships vary across stimulus types (Fig. 4m, see also Fig. S4-1j,k). We find that at least 35% of cluster assignments are retained across different stimuli, compared with 60-75% cross-validation within a single stimulus condition. Clusters obtained from the looming stimulus are the most distinct from other stimuli, which is consistent with the earlier observations from regression analysis (Fig. 2e-h). Interestingly, clusters identified from the spontaneous period (featureless environment) match significantly (≥50%) with clusters obtained from phototactic stimuli, moving stripes (OMR) or dark flashes conditions. These cross-validation results suggest that the organization of functional clusters can be variable and context-dependent.

The regression and clustering approaches can be flexibly and interactively combined with anatomical knowledge to identify and investigate neural circuits (Fig. S4-2a). We found a rich variety of circuits, some of which span large distances across the brain, thanks to the whole-brain coverage of our data. We have included a few example circuits to demonstrate the effectiveness of this integrated approach (see Supplemental Text accompanying Fig. S4-2), including a jaw and gill movement control circuit (Fig. S4-2b), mesencephalic locomotion-related region (Fig. S4-2c), the raphe nucleus and the vagus cranial system (Fig. S4-2d,e), the olfactory bulb (Fig. S4-2f), and a putative eye movement control circuit that we describe in detail below.

### Abducens nucleus / eye movement control

Previous studies (Miri et al., 2011a, 2011b; Portugues et al., 2014) have located eye-movement related neurons in the hindbrain by correlating neural activity to eye-tracking data, but clear consensus is lacking about the precise location of this neuronal population in zebrafish. Although we did not have behavioral recordings of eye position, we could nonetheless identify clusters exhibiting functional and anatomical features that suggest these clusters are related to eye-movement control. Two pairs of anatomically compact clusters in the ventral hindbrain in rhombomeres 5 and 6 consistently appear in the whole-brain clustering for most fish, and can be manually identified in all fish by clustering neurons in the corresponding locations (Fig. 4n, arrows with tails; most caudal cluster corresponds to Z-Brain mask “6.7FDhcrtR-Gal4 Cluster 3”). The cells in these clusters stand out because their activity traces are highly correlated to each other (many correlations exceed 0.8, see Fig. S4-1o), but not significantly correlated to either the stimulus or the fictive swimming behavior (Fig. 4r). The anatomy of these clusters closely matches that of the abducens nucleus (ABD) characterized in goldfish, which consist of motor neurons (arranged in two nuclei in rhombomeres 5 and 6) that are electrically coupled by gap junctions (Cabrera et al., 1992; Gestrin and Sterling, 1977). Indeed, the high correlations we observe within these clusters may be caused by gap junction connections.

We then screened for putatively eye-movement related neurons by regressing to the cluster center of the ABD clusters (Fig. 4n). This analysis is analogous to Fig. 2c, except here the ABD activity is used as a proxy for motor output (i.e. eye movements). This revealed an additional pair of small clusters near the anterior border of the hindbrain that are tightly correlated to contralateral ABD clusters, which we identified as the oculomotor nuclei (OCM) based on anatomical labels in the Z-brain atlas (Fig. 4n, arrowheads). In the goldfish, two groups of interneurons that are located in close proximity to the ABD nucleus project contralaterally to the OCM; thus close functional relationships between the ABD nuclei and contralateral OCM clusters are expected (Cabrera et al., 1992; Gestrin and Sterling, 1977). To characterize the sensory-motor organization of this circuit in more detail, we mapped the anatomical locations of highly motor (ABD) related cells, colored according to their functional activity in a sensory-motor space (Fig. 4p, amplitude of periodic component vs regression coefficient with left motor res., Fig. 4q, anatomical map, see Fig. S4-2g for corresponding figures for right motor res.). The two putative ABD clusters in the ventral hindbrain (rhombomeres 5 and 6) appear to have a core/shell anatomical structure: cells on the dorsal and anterior edges of the clusters are less motor correlated and more similar in activity to the OCM clusters. We also observed neurons in the more dorsal sections of the hindbrain with reduced motor correlation and enhanced sensory components. We speculate that these neurons could be responsible for the integration of sensory inputs for eye-movement control. Based on this observation and studies of the circuit controlling eye-movement in the literature, we propose a diagram of eye-movement circuit with components consisting of the populations of neurons identified above (Fig. 4o). Using a combination of regression and clustering analyses on whole-brain activity data, we have been able to postulate a connectivity diagram for a circuit for which we have neither the motor output (eye-tracking) nor a robust stimulus drive (stimuli used here do not strongly drive eye-movement). This demonstrates the power of the whole-brain data and in-depth analysis for generating circuit hypotheses that invite detailed follow-up studies.

### Multi-stimulus convergence

In general, distinct sensorimotor pathways that culminate in the same behavioral output are expected to converge to a common motor pathway. In our experiments, different visual paradigms eventually converge at the motor system to generate swimming behavior (i.e. left and right turns). Here we asked where the first site of convergence between the phototaxis (phT) and ocular-motor response (OMR) sensorimotor pathways is located, defining convergent activity as scoring highly in regression to both phT and OMR stimuli that drive left turns, and likewise for right turns. A map of highly convergent cells (top 3%, Fig. 5a, see also Fig. 2e,f and Fig. S5b) reveals well-defined groups of neurons in the anterior hindbrain, specifically in Rhombomere 1 and the medial area of Rhombomere 2. This anatomical region is composed of several clusters with distinct functional activity, as revealed by whole-brain clustering (Fig. S5e). Interestingly, these clusters form medial/lateral stripes that may correspond to the neurotransmitter stripes in the ARTR (see also Fig. S5a) (Dunn et al., 2016b).

**Figure 5.**
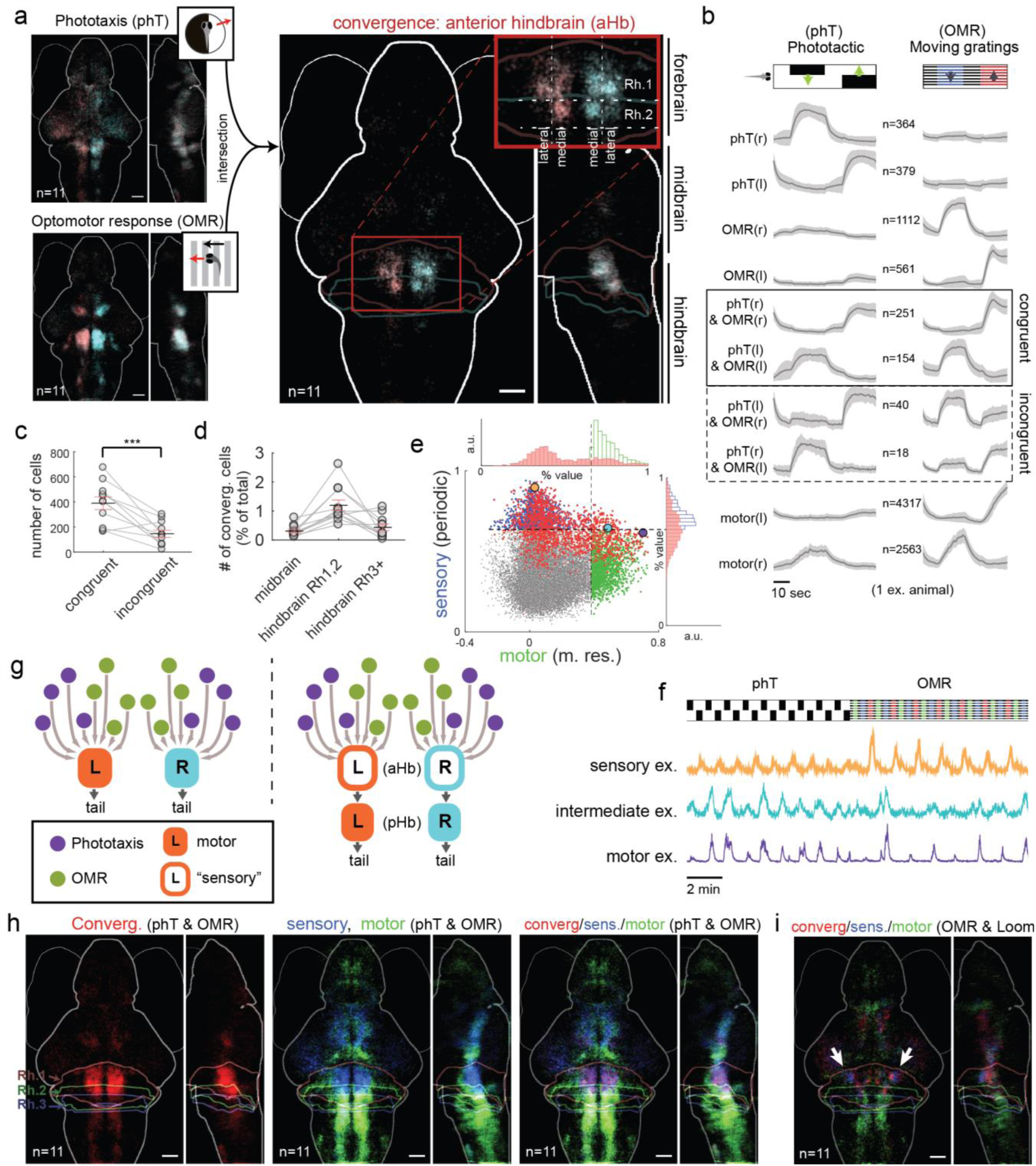
Multi-stimulus integration. (**a**) Average map across fish for the intersection of phototactic responsive cells and OMR responsive cells. Multi-stimulus responsive cells are concentrated in Rh.1 and the medial stripes of Rh.2. (**b**) Whole-brain regressions are performed to a set of regressors that include phT only, OMR only, phT&OMR joint (including congruent and incongruent), and motor output regressors. Cells that have a correlation coefficient >0.4 to at least one of these regressors are classified by their best regressor. Average functional activity of cells associated with each regressor is plotted (dF/F, mean±SD). (**c**) Quantification of the number of cells with activity classified as congruent or incongruent (n = 18 fish). Congruently tuned cells significantly outnumber incongruently tuned cells. (**d**) Quantification of the number of convergent cells (normalized as fraction of total cells) for midbrain, hindbrain Rh1&2, and hindbrainRh3+ regions. Anterior hindbrain Rh1,2 contains the largest fraction of convergent cells. (**e**) Scatter plot of convergent activity in a 2-dimensional sensory-motor space. Each point represents a cell. Horizontal axis; motor component: regression coefficient of cellular activity with left motor res. (see Fig. 3d). The maximum value from phT or OMR blocks is plotted. Vertical axis: sensory component: square root of the variance explained by periodic component of activity. The minimum value from phT or OMR blocks is plotted. Red points: convergent cells, defined as the intersection of the top 5% of cells ranked by phT and OMR regression (as in Fig. 5a, but with 5% threshold). Blue points: top cells ranked by sensory component (same number of points as red). Green points: top cells ranked by motor component (same number of cells as red). Top and right: histograms of motor and sensory components, comparing distribution of convergent cells (red) to the most motor or sensory-related cells, respectively. (**f**) Cellular activity for 3 example neurons shown in (e) during phT and OMR stimulus blocks. (**g**) Illustration of two alternative hypotheses for multi-sensory integration. *Left:* information from non-overlapping visual representations (e.g. phT, OMR) directly feeds into premotor systems, which then compete to produce different behaviors. *Right:* different visual representations first feed into a behavior-centric visual representation before affecting motor circuits. The present results support the second model, with the anterior hindbrain containing the behavior-centric, convergent visual representations. (**h**) Average anatomical maps (n = 11 fish) showing location of the cells represented in (e): top-ranking convergent cells (left map, red), sensory-related cells (middle map, blue) and motor-related cells (middle map, green), and merge (right map, overlap between sensory-related and convergent appears purple. Note significant overlap in the anterior hindbrain. Boundaries for Rhombomeres 1,2, and 3 are overlaid. Scale bars, 50 µm. (**i**) Same as (h), but for convergence between OMR and looming stimuli.

With two types of stimuli, there are several types of responses we might expect to see, including cells corresponding to phT-only, OMR-only, both phT and OMR, and motor (Fig. 5b, Fig. S5h,i). It is also in principle possible for cells to respond to incongruent stimuli pairs, i.e. phT-left and OMR-right, or phT-right and OMR-left. Indeed, using regression, cells can be found that respond to both congruent (convergence cells) and incongruent stimuli pairs. However, the number of congruent cells is much greater than incongruent (Fig. 5c) and congruent cells are most numerous in the anterior hindbrain (Fig. 5d, Fig. S5c,d,g). Since the distinction between congruent and incongruent stimuli is defined by motor output, these observations suggest that the convergent activity in the anterior hindbrain plays an important role in the sensorimotor transformation.

Because the behavioral response is similar for congruent phT and OMR stimuli, motor-related activity will by definition be somewhat convergent. However, motor-related activity is conceptually distinct from **sensory convergent** activity, which we define as being tuned primarily to congruent stimuli and not directly related to motor output. Thus, cells labeled as convergent (Fig. 5a) can be either sensory convergent or motor-related. The distinction between sensory convergent and motor-related activity can be understood via two competing circuit models. In a direct motor convergence model, sensory activity for phT and OMR converge directly on motor-related cell clusters (Fig. 5g, left). In a sensory convergence model, sensory activity first converges in sensory convergence clusters, which in turn activate the motor-related clusters (Fig. 5g, right).

To distinguish the two models, we leveraged analysis of trial-to-trial variations (used similarly in Fig. 3). We would expect sensory convergent cells to exhibit concerted periodic activity during the presentation of both phT and OMR stimuli and lack aperiodic, motor-related activity. Conversely, we expect motor-related cells exhibit aperiodic motor-related activity and lack periodic activity. To differentiate these two types of responses, we plotted cell activity of convergent sensory cells (top 5% for both regressions, similar to Fig. 5a) in a two dimensional sensory-motor space (Fig. 5e, red points) where the y axis quantifies sensory-related periodic activity and the x axis quantifies aperiodic motor activity (see legend for details). To facilitate comparisons, we also plotted all neurons (gray points), and highlighted the most sensory (blue points) and most motor (green points) cells, each matched in number to the convergent cells (red). Among the convergent cells, we observe strongly sensory convergent cells (Fig. 5e,f, orange dot and trace) and strongly motor-related cells (Fig. 5e,f, purple dot and trace), as well as cells intermediate between the two (Fig. 5e,f, cyan dot and trace). The distribution of sensory convergent, intermediate, and motor-related cells is evident from the sensory-motor scatter plot (Fig. 5e, see also corresponding histograms top and right), with sensory convergent cells occupying the top left part of the plot, motor-related on the right, and intermediate cells in between. The distribution of cells in this sensory-motor space varies somewhat across different fish and turning directions (see Fig. S5l for corresponding right-turn activity), but in all fish a significant portion of sensory convergent cells overlap strongly with the most sensory cells (Fig. S5j,k).

We also examined the anatomical distribution of convergent cells (Fig. 5h, left, multifish average), and compared them to the anatomical distribution of the most sensory-related cells and most motor-related cells (out of all cells) (Fig. 5h, middle; cells selected as in Fig. 5e). There is considerable overlap between convergent cells and sensory-related cells in rhombomeres 1 and 2 in the hindbrain (Fig. 5h, right, purple cells), and much less overlap between convergent cells and motor-related cells in the more posterior hindbrain (rhombomeres 3 and higher). These maps are consistent with the hypothesis that there exists a population of sensory convergence cells that reside in the anterior hindbrain, while motor-related cells tend to be located in more posterior segments. A similar analysis for a different pair of stimuli, namely OMR and looming, also identifies sensory convergence populations (purple) in similar areas of the anterior hindbrain (Fig. 5i, arrows), but restricted to more lateral locations compared to the phT/OMR sensory convergence cells.

We hypothesize that the existence of a significant population of sensory convergence neurons is an important step in the generalized sensorimotor transformation. The identification of such a population disproves the direct motor convergence model (Fig. 5g, left), and supports a sensory convergence model (Fig. 5g, right). The anatomical location of the sensory convergence cells, bordering both the midbrain (where neurons processing visual features reside) and motor nuclei such as the ARTR, makes it convenient to form synaptic connections with both. While additional experiments will be required to understand in more detail the role played by this sensory convergence area, our whole-brain imaging experiments have constrained possible circuit models and can precisely guide future investigations.

## Discussion

In this study, we generated a multi-animal, whole-brain, cellular-resolution dataset incorporating multiple stimulus paradigms and developed extensive analyses aimed at understanding the functional organization of the larval zebrafish brain. We mapped out the sensory and motor components of brain-wide brain activity and dissected the neural correlates of behavior in detail. To further extend the analysis beyond sensory and motor-related neural dynamics, we developed a customized functional clustering method to reveal a comprehensive collection of representative neural activity profiles. Using a combination of analysis techniques, we identified and examined a circuit putatively involved in eye-movement control. Lastly, we identified anatomically well-structured functional groups in the anterior hindbrain that generalize stimuli that elicit similar turning responses.

We started out with a relatively straight-forward regression analysis identifying cells tuned to specific stimuli or motor output (Fig. 2). We found that there is a relatively large and dense population of motor-related neurons in the hindbrain that also exhibit highly concerted activity, which may be somewhat redundant from the view-point of coding capacity. Interestingly, the hindbrain segments containing the most concerted activity occupy the same Rhombomere sections as the mammalian pons, and contain many homologous nuclei (Kandel et al., 2013). We speculate that evolutionarily older brain areas may generally rely on more concerted activity. Alternatively, highly concerted activity may also be a feature of a developmentally young brain, and motor activity may differentiate further in adult fish to allow for more nuanced motor control.

Having identified cells related to sensory stimuli and motor outputs, we then sought to understand the sensorimotor transformation in more detail. The conceptual framework of the sensorimotor transformation is easy to oversimplify. Intuitively, it is tempting to think of the sensorimotor transformation as a one-dimensional axis in which the sensory input gradually morphs into motor output. However, in this framework, sensory and motor-related activity cannot be distinguished, because a motor neuron that is driven strongly by stimulus would be highly correlated to stimulus regressors as well. This one-dimensional model also fails to account for variability in the behavior (i.e. motor signals that occur unrelated to the stimulus). Motor variability thus plays a key role in expanding the sensorimotor framework (Renart and Machens, 2014). In our analysis, we extracted the trial-to-trial variation of the motor output to differentiate “purely motor” activity from sensory-related or sensory-driven components. We plotted anatomical tuning maps for the separate trial-average and residual motor components (Fig. 3f), and found that cells related to motor variability are broadly distributed in the hindbrain exclusively posterior to the Rhombomere 2/3 boundary (Fig 3f, lower right). Interestingly, the distribution of cells related to motor variability was largely overlapping with the distribution of all motor-related cells (Fig. 2c), and we did not observe any particular anatomical location exclusively correlated to motor variability. This suggests that motor variability does not come from a particular anatomical nucleus, but may instead depend on many contributing factors, such as neuromodulatory systems or internal states. It is also possible that behavioral variability originates as small perturbations that are amplified by the motor systems.

We identified cells whose activity is highly correlated to both trial-average and residual motor activity, postulating that they play a key role in the sensorimotor transformation (Fig. 3f, upper right). However, a caveat is that correlation-based analysis cannot distinguish feedforward versus feedback motor-related activities. Determining the directionality of functional connections would probably require controlled perturbation experiments, although specialized computational analysis paired with high-framerate neural activity recordings may also be productive. One promising target for future studies would be to relate the functional relations within and between clusters to a systematic dataset of anatomical connectivity in zebrafish (Hildebrand et al., 2017).

It is important to note that the fast, near-simultaneous nature of light-sheet imaging is crucial to our ability to dissect independent sensory and motor residual components. The residual (non-trial-average) activity of cells can only meaningfully be compared if activity of all cells is recorded nearly simultaneously, which requires the fast frame rates (∼2-3 volumes/sec) characteristic of light-sheet data. Near-simultaneous data collection is also essential for the functional clustering analysis, which allowed us to identify concerted activity that is not strongly correlated to either sensory or motor events and requires concurrent observations of neuron groups in faraway brain regions.

Our clustering algorithm is a variation of agglomerative clustering with additional specific features tailored to our data (see Methods) that endow our algorithm with robustness to noise and high computational efficiency; whole-brain single-cell resolution data (∼100,000 cells, ∼5000 time frames) can be efficiently clustered on a standard desktop computer on the timescale of minutes. The algorithm is sensitive enough to detect even very weak functional clustering patterns in the data, including artefactual signals resulting from the scanning laser itself (Fig. S4-1i, these artefactual clusters were thereafter excluded from analysis).

In addition to identifying specific clusters, whole-brain automatic clustering analysis reveals three broad characteristics of the functional organization of the larval zebrafish brain. First, a significant percentage of neurons do not correlate strongly to any clusters (the unclustered neurons in Fig. 4c, see also Fig. S4e for varying correlation threshold), as has also been observed in other neural systems (Okun et al., 2015). Many of these functionally isolated neurons may be still be developmentally immature (Boulanger-Weill et al., 2017), as they are especially common in areas where young neurons are added to the rapidly developing brain (see Fig. S3c). Others may be computationally important despite remaining unidentified by clustering, and understanding their functional roles may require more complex analysis or perturbation experiments. Second, functional activity among different clusters varies gradually, and most clusters exhibit significant correlations with other clusters (Fig. 4i). Moreover, most characteristics of the clustering, including number of clusters, cross-validation score, and total number of cells included in clusters vary continuously with the clustering threshold (Fig. S4-1d). This suggests that the functional relationship between clusters depends on the chosen correlation scale, and that there is no clear correlation scale where brain-wide organization is most evident. As a result, the relationships between functional clusters are complex and likely best described hierarchically (Fig. S4-1f). Third, the extent to which functional clusters are anatomically conserved varies across the brain (Fig. S4-1h); this is consistent with previous observations of the positioning of transgenic labels (Randlett et al., 2015). It still remains to be determined what factors are responsible for this observed stereotypy/variability of the functional organization. An intriguing hypothesis is that some, but not all, regions of the brain are self-organized through experience-dependent plasticity, which may lead to across-animal variability on smaller anatomical scales.

Starting with the experimental design incorporating different visual stimuli, we set out to understand in more detail how sensory representations converge before motor output. The overlap of different stimulus tuning maps (Fig. 2e-h) in the anterior hindbrain reveals that this area is generally important for sensory processing, and the concentration of convergent congruent activity in the same area (and relative lack of incongruent tuning) makes it a candidate circuit for informing behavior based on diverse sensory input. The distinction between the observed convergent sensory representation and more motor-related representations indicates that sensory convergence is at least a two-step process – first sensory convergence, then motor output (Fig. 5g, right model as opposed to left). However, it is important to realize that cellular responses form a continuum (Fig. 5e), and there exists a gradient between the purely sensory and and purely motor-related clusters in the hindbrain. The sensory convergent cells contain an abstraction of the stimulus, encoding “right-turn-eliciting stimuli” or “left-turn-eliciting stimuli”, even in cases where the matching motor output is absent. This generalization of congruent sensory stimuli *before* the motor output ultimately may allow for a much more flexible and complex sensorimotor transformation.

Based on our observations that sensory convergent cells are tuned to different congruent stimuli, it is reasonable to hypothesize that these cells may also integrate multiple simultaneously presented stimuli. In this context, the activity of the sensory convergent cells may encode the overall influence of the stimuli on the motor output. In cases where competing incongruent sensory stimuli are presented, such a role could be important for a type of decision-making. While our results based on serially presented stimuli do not directly address this hypothesis, they nevertheless suggest that the sensory convergent cells in the aHB may be a hub for decision-making computations.

We have shown that in-depth explorations of functional whole-brain activity data can generate strong hypotheses about sensorimotor processing and the broader functional organization of the brain. The analytical methods presented here are also more generally applicable to other large-scale, high-resolution functional datasets. We have made our software platform and data openly available to facilitate further analysis and adaptation of this platform for other experimental studies. The analysis of the functional organization in brain-wide circuits is complementary to techniques of optical manipulation, electrophysiology, viral tracing, and connectomics, and in combination with these techniques, offers the potential to promote our mechanistic understanding of brain-wide circuits.

## Acknowledgements

We would like to acknowledge Takashi Kawashima for help with the preprocessing analysis, and members of the Engert lab for helpful discussions. This work was supported by the Howard Hughes Medical Institute (M.B.A. and Y.M.), the Simons Collaboration on the Global Brain (awards #325171: Y.M. & M.B.A, #542973: M.B.A. and F.E., and #325207: H.S.), NIH grants from the NINDS (1U19NS104653: F.E. and H.S. and U01NS090449: F.E.), and the Gatsby Charitable Foundation (H.S.).

## Author contributions

Conceptualization, X.C., M.B.A., F.E., Y.M., Y.H., and H.S.

Methodology, X.C., Y.M., Y.H., A.T.K., and M.B.A.

Software, X.C.

Formal Analysis, Y.H., X.C., A.T.K., and Y.M.

Investigation, Y.M.

Resources, M.B.A., F.E., and H.S.

Data Curation, X.C., Y.M., O.R. and M.N.

Writing – Original Draft, X.C., A.T.K., Y.H., Y.M., M.B.A., H.S., F.E.

Writing – Review & Editing, X.C., A.T.K., Y.H., Y.M., M.B.A., H.S., F.E.

Visualization, X.C.

Supervision, F.E., M.B.A. and H.S.

Funding Acquisition, F.E., M.B.A. and H.S.

## STAR Methods Text

### Contact for Reagent and Resource Sharing

Further information and requests for resources and reagents should be directed to and will be fulfilled by the Lead Contact, Xiuye Chen.

### Experimental Model and Subject Details

Transgenic zebrafish, panneuronally expressing calcium indicator GCaMP6 under the *elavl3* promoter and nucleus-targeted as *Tg(elavl3:H2B-GCaMP6f)*, was used for imaging (Vladimirov et al., 2014). Zebrafish larvae (5 dpf – 7 dpf), were paralyzed with 1mg/ml alpha-bungarotoxin (Sigma-Aldrich), and embedded with 2% low melting point agarose. All experiments presented in this study were conducted in accordance with the animal research guidelines from the National Institutes of Health and were approved by the Institutional Animal Care and Use Committee and Institutional Biosafety Committee of Janelia Research Campus.

### Method Details

#### Light-sheet imaging

Light-sheet imaging experiments were performed with an experimental setup previously described (Vladimirov et al., 2014), which achieves almost whole-brain imaging with concurrent presentation of visual stimuli and electrical recordings of fictive swimming. The imaging rate was about 2 brain volumes/s (2.11 ± 0.26 Hz), with the variability due to differences in the size of the brain for different animals (mainly because of the thickness difference). Each experiment lasted between 30 and 120 min.

#### Cell detection

After imaging acquisition, the time series of each plane was registered by custom-written C/CUDA software for the XY plane translation. Z-drift was corrected by first comparing patches of images with the nearby planes from the first 2 min of the recording, and then calculated by linear fitting the distance across the Z-planes. Only the parts of experiment when the XY- or Z-drift smaller than 1 um was used for further analysis.

Cells were detected from the time-averaged image. In the *Tg(elavl3:H2B-GCaMP6f)* line, calcium indicator was mainly localized in the nucleus and forms a bright disk on each plane, a property that facilitates neuron detection. First, GCaMP expression area was extracted by binary thresholding based on pixel intensity and local contrast. Second, each pixel was normalized locally by assigning a relative rank of intensity within a disk patch (radius = 4 µm), then further smoothed by a circular patch with radius of 1.6 µm. The center of a cell body was identified as being a local maximum point, and the calcium trace for the cell was calculated as the average over circular patch with radius 2.8 µm (Kawashima et al., 2016).

#### Fictive behavior recording

The fictive behavior setup has been previously described (Dunn et al., 2016b). Two suction glass pipettes (∼45 µm inner diameter) were attached on the skin from each side of the tail. Gentle suction was applied to help the electrical contact with the motor neuron axons. These electrodes record spiking from multiple motor neuron axons, providing readout of intended locomotion (Ahrens et al., 2012; Dunn et al., 2016b; Masino and Fetcho, 2005). Extracellular signals were amplified (Molecular Devices, Axon Multiclamp 700B), fed into a computer using a National Instruments data acquisition card, and recorded by custom written C# software. The fictive swim bouts were first detected as previously described (Ahrens et al., 2012, 2013), then used to decode fictive turns (Dunn et al., 2016b). The extracellular signal from left and right side of the tail were recorded from two independent channels of the amplifier. The fictive swim signals were calculated as the smoothed power of the deviation from baseline. Individual swim bouts were detected automatically, and then a weighted average swim bout amplitude was used to normalize and balance the signal from left and right channel. Each swim bout was weighted by a normalized rising exponential function, to take into account the fact that turns affect the start of swim bouts more heavily than the end of swim bouts. For the same reason, to determine the fictive turn amplitude and distance, filtered and normalized fictive signals during swim bouts were weighted with a decaying exponential function (tau = [bout duration]/3) to emphasize the initial burst that determines overall turn direction. Then the turn amplitude was calculated by the difference of weighted power divided by the sum of the weighted power from both sides: (Power_Left_ - Power_Right_) / (Power_Left_ + Power_Right_). The sum of the weighted power was used to measure the swim distance.

#### Visual stimulation

Visual stimulation patterns were generated by custom-written C# software, delivered by a projector with homogenous red light, and projected to a diffuser that was stuck to the bottom of the imaging chamber. The visual stimulation consisted of serial repetitions of different sets of patterns: For **phototaxis**, two patterns were presented: Left-Dark / Right-Bright, and Left-Bright / Right-Dark. After each pattern a period of whole field dark was presented. The dark area slightly crossed the midline in order to motivate the fish to swim towards the bright side. For **moving grating** stimulation, three or four directional drifting gratings were presented: forward, backward, left-ward, and right-ward. Forward and backward gratings moved along the body axis towards the head or the tail direction. Left- and right-ward gratings were oriented 120 degrees away from the head direction, either to the left or the right side. Grating speeds were as follows: Leftwards, rightwards, backwards: 0.4mm/sec. 6 degree/sec. Forwards: 1 mm/sec, 15 degree/sec. For **spontaneous** stimulation, the whole field was dark and no other visual feature was presented. For **dark flash** stimulation, whole field dark and whole field light were presented alternatingly for 20 sec each. For looming stimulation, a bright background was presented, and a dot was expanded on either left or right side of the fish. Expansion lasted for 5 seconds, and was followed by 25 seconds of whole field bright.

#### Image Registration

Nonrigid image registration was done with CMTK (http://www.nitrc.org/projects/cmtk/), similar to registration done for the Z-Brain atlas (Randlett et al. 2015). Additionally, one “bridge brain” was created by imaging the same fish both with the light-sheet microscope (used for the functional dataset in this study) and a confocal microscope (used for the creation of the Z-Brain atlas). Functional light-sheet datasets were first registered to the bridge brain, and the transformations from light-sheet to confocal for this bridge brain were subsequently applied to complete the registration. To obtain the anatomical location of individual cell-ROI’s in the transformed coordinates, we used the CMTK built-in command “streamxform”.

#### MATLAB GUI

A graphical user interface (GUI) was written in MATLAB® for interactive data visualization and analysis. For each fish, the functional data loaded consisted of one calcium trace per each segmented cell, calculated as change in fluorescence (ΔF/F). Also loaded were the annotated stimuli, fictive behavior, anatomical location of segmented cells (both raw and registered), anatomy stacks, and annotated Z-Brain masks. The GUI integrates most analyses used in this study, including but not limited to manual cluster selection, selection based on anatomy, set operations, regression methods, unsupervised clustering methods, storage of clusters, integration with the Z-Brain atlas, and various visualization and export options. The software was written and tested with MATLAB 2016a and running on Windows 7.

#### Automated Functional Clustering algorithm

This algorithm was custom developed to suit this dataset, and the code is available as part of the GUI. We outline the algorithm below:

1. Divide all cells into “functional voxels” (∼10 cells each)

a. Perform k-means clustering on all cells (k=20)
b. Perform k-means clustering on outputs of (a) (k = ∼400)
c. Discard any cells whose correlation with the voxel average activity is less than $THRESH
d. Discard any voxels with fewer than 5 cells
2. Merge voxels into clusters based on density in functional space

a. for each pair of voxels ij (starting from most correlated): if the correlation between voxel i and j is greater than $THRESH, and the correlation between the the voxel j and the centroid (average) of the cluster containing voxel i is greater than $THRESH: then group voxel j in the same cluster as i.
b. discard any clusters with fewer than 10 cells
3. Clean up clusters using regression to cluster centroids

a. for each cell k: if the correlation to the closest cluster’s centroid is greater than $THRESH: include cell k in that cluster
b. Discard any clusters with fewer than 10 cells
4. Iterate merge and cleanup steps

a. Perform step 2 and 3 once more, using clusters as input voxels.

This clustering algorithm can either be applied to all cells in the brain or a chosen subset of interest, and the correlation threshold determining clustering stringency ($THRESH) can be adjusted to trade-off completeness and accuracy (see Results and Fig. S4-1d). For most analysis in the text, the value of $THRESH was 0.7.

#### Clustering Cross-Validation

We divided the data into two halves along the time dimension; where multiple stimulus repetitions were present, we used an equal number of repetitions for the two halves. We then use the Hungarian method (Munkres assignment algorithm) to match clusters produced from the two halves of the data based on the fraction of common cells contained in a pair of clusters. This matching index was also visualized as the total “mass” distributed along the diagonal entries (Fig. S4-1e). The cross-validation coefficient (Fig. 4m) is the fraction of cells that were assigned membership to the same cluster for both halves of the data.

#### Screen for anatomically conserved clusters

We used the following criteria to determine whether two clusters were considered occupying comparable locations in anatomical space. First, given a pair of clusters, we calculated the distance of all pairs of cells within each cluster (sets *d*_*1*_ and *d*_*2*_), and the distance of all pairs of cells between the two clusters (*d*_*12*_). If the relative distance mean(*d*_*12*_) / min(mean(*d*_*1*_), mean(*d*_*2*_)) was less than 2, we considered the clusters conserved (with some restrictions on cluster sizes to exclude outliers). For each cluster in a given fish, we compared it with all clusters in all other fish. If there were at least 6 other fish that contained at least 1 matched cluster based on the criteria above (multiple matching clusters were possible), this cluster was marked as a conserved cluster over the population (Fig. 4l).

#### Stimulus and motor regressors

The set of stimulus regressors that represent different stimulus features (Fig. 2d-h) were generated by convolving box-car regressors with a single exponential kernel for the calcium indicator GCaMP6s (half-time 0.4 sec, peak delay 0.08 sec).

Motor regressors were generated from recorded fictive behavior (forward, leftward and rightward swimming) convolved with the temporal filter of the calcium indicator (exponential function, half-time 0.4 sec, peak delay 0.08 sec). Though sufficient for some analyses (Fig. 2c), this simple convolution does not fully account for the relation between behavior and the nuclear-labeled calcium trace. In particular, the calcium traces of neurons in the motor region have a longer time constant than the convolved behavior. For some regression maps (Fig. 3d,f and Fig. 5e,h), we found that the results can be improved by using “motor seed” traces as the behavior regressor. The motor seeds are a small number of neurons manually selected in each fish based on two criteria: (i) the neurons have the highest correlations to the fictive behavior and (ii) they are located in the region of the hindbrain that is known to send output signals to drive swimming behavior. The average activity traces of the motor seed cells were denoted “motor outputs” in the manuscript (as opposed to “motor regressors”). In analysis for Fig. 2, the fictive motor regressors were used, whereas in Fig. 3 and Fig. 5, the motor outputs taken from motor seeds were used.

#### Decomposing activity into trial averages and trial residuals

For any trace, we index its value at trial k and time t (relative to the beginning of the trial) as *x*_*t,k*_. Its trial average component is defined as 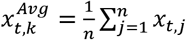 (n is the number of trials), and its trial residual component is 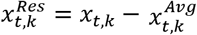. Importantly, *x*^*Avg*^ and *x*^*Res*^ are orthorgonal, because 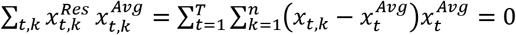 (T is the period of the stimulus). In fact, *x*^*Res*^ is orthogonal to any trace that is periodic to the stimulus, according to the same argument. In particular, let S represent the subspace of all periodic traces locked to stimulus trials, then *x*^*Avg*^ is the orthogonal projection of x in S.

The sensory periodicity (as in Fig. 3f and Fig. 5e) is calculated as the square root of the fraction of the variance of *x*^*Avg*^ relative to x. The square root is taken so that the resulting index is comparable in scale to correlation coefficients (e.g. consider the correlation between the trial-average and the original trace) that we use in other analyses such as the correlation to motor-residuals and motor-averages.

When applying the decomposition into trial-average and trial-residual to motor traces, the orthogonality of the decomposition removes the correlation between motor and stimulus, and regressing to motor_res hence improves the identification of motor related neurons as used in Fig. 3 and 5.

As shown above, the motor-res is orthogonal to S. This means that (correlation to motor_res)^2^ + (sensory periodicity)^2^ = the fraction of variance of *x* within the joint subspace spanned by {motor-res, *S*} ≤ 1. Note that the subspace spanned by {motor-res, S} is the same as that spanned by {motor, S}. So the amount of remaining activity 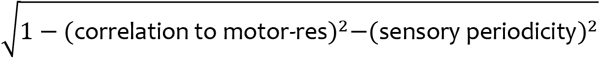 represents activity beyond sensory and recorded motor behavior.

#### Quantification and Statistical Analysis

Fig. 2a, bottom: mean±SD dF/F plotted for cells with regression coefficient > 0.5

Fig. 2b: shuffle traces obtained by randomizing time indices for activity traces

Fig. 2c: cells with regression coefficient > 0.5 plotted, left: single fish, right: n=16 fish superimposed

Fig. 2d: mean±SD dF/F plotted for cells with regression coefficient > 0.5

Fig. 2e: cells with regression coefficient > 0.5 plotted, n=7 fish superimposed

Fig. 2f: cells with regression coefficient > 0.5 plotted, n=11 fish superimposed

Fig. 2g: cells with regression coefficient > 0.5 plotted, n=9 fish superimposed

Fig. 2h: cells with regression coefficient > 0.5 plotted, n=11 fish superimposed

Fig. 3b: mean±SD dF/F plotted for cells within curated k-means clusters

Fig. 3f, maps: top 2% of cells by regression rankings plotted for each fish, n=17 fish superimposed

Fig. 3f, scatter plot: each cell represented by a point. x value is regression coefficient for motor res. for that cell. y value is regression coefficient for motor avg. for that cell. Data for a single example fish is shown, with motor regressors derived from the left-side motor output. See methods for definition of motor output, motor avg. and motor res.

Fig. 3g, maps: top 2% of cells by regression rankings plotted for each fish, n=17 fish superimposed

Fig. 4c: t-SNE for one example fish only

Fig. 4d: Histogram of number of cells included in each cluster, mean±SEM across n=18 fish plotted for each bin

Fig. 4e: Histogram for average correlation between cells for each cluster, mean±SEM across n=18 fish plotted for each bin

Fig. 4f: Histogram of average anatomical distance between cells for each cluster, mean±SEM across n=18 fish plotted for each bin

Fig. 4i: Matrix of Pearson correlation coefficients between pairs of clusters.

Fig. 4l: Map of clusters that are conserved for at least n=6 fish. Conserved clusters are superimposed for a pool of n=18 fish. See methods for details of definition of “conserved”.

Fig. 4m: Two-fold cross-validation showing fraction of clusters that pass a cross-validation test. See Methods for details of cross-validation. Average fraction of clusters shown for n=6 fish.

Fig. 4n: Map of ABD clusters, n=17 fish superimposed.

Fig. 4p: Each cell represented by a point. x value: square root of the variance explained by the periodic component of the cell’s activity. y value: regression coefficient for motor res. for that cell.

Fig. 4r: Average activity of putative ABD nuclei for several fish. Each pair of red/blue traces show average left/right ABD nuclei activity for one fish.

Fig. 5a: phT and OMR maps show top 3% of cells by regression coefficient ranking. Convergence map shows cells included in both phT and OMR maps.

Fig 5b: Cells that have a correlation coefficient > 0.4 to at least one of the regressors shown are classified by their best regressor. mean±SD dF/F plotted for cells all taken from the same fish. n refers to the number of cells included in each category.

Fig. 5c: Number of congruent and incongruent cells (as defined in Fig. 5b) for n=18 fish. mean±SEM shown in red. p<.001, Student’s t-test.

Fig. 5d: Number of convergent cells (defined in Fig. 5a) found in different brain areas for n=18 fish. mean±SEM shown in red.

Fig. 5e: scatter plot: each cell represented by a point. x value: regression coefficient for motor res. for that cell. y value: square root of the variance explained by the periodic component of the cell’s activity. Data for a single example fish is shown, with motor regressors derived from the left-side motor output. Red points: convergent cells, defined as intersection of top 5% phT and OMR regression maps. Blue points: top cells ranked by y value, same number as red cells. Green points: top cells ranked by x value, same number as red cells. Top: histogram of x values for red and green cells. Right: histogram of y values for red and blue cells.

Fig. 5h: Maps of red, green and blue cells (defined as in Fig. 5e), superimposed for n = 11 fish.

Fig. 5i. Similar to Fig. 5h, but for OMR and looming convergence, rather than phT and OMR.

#### Data and Software Availability

Software and instructions for downloading the data will be made available after peer review of this manuscript.

## Supplemental Material

**Figure S1.**
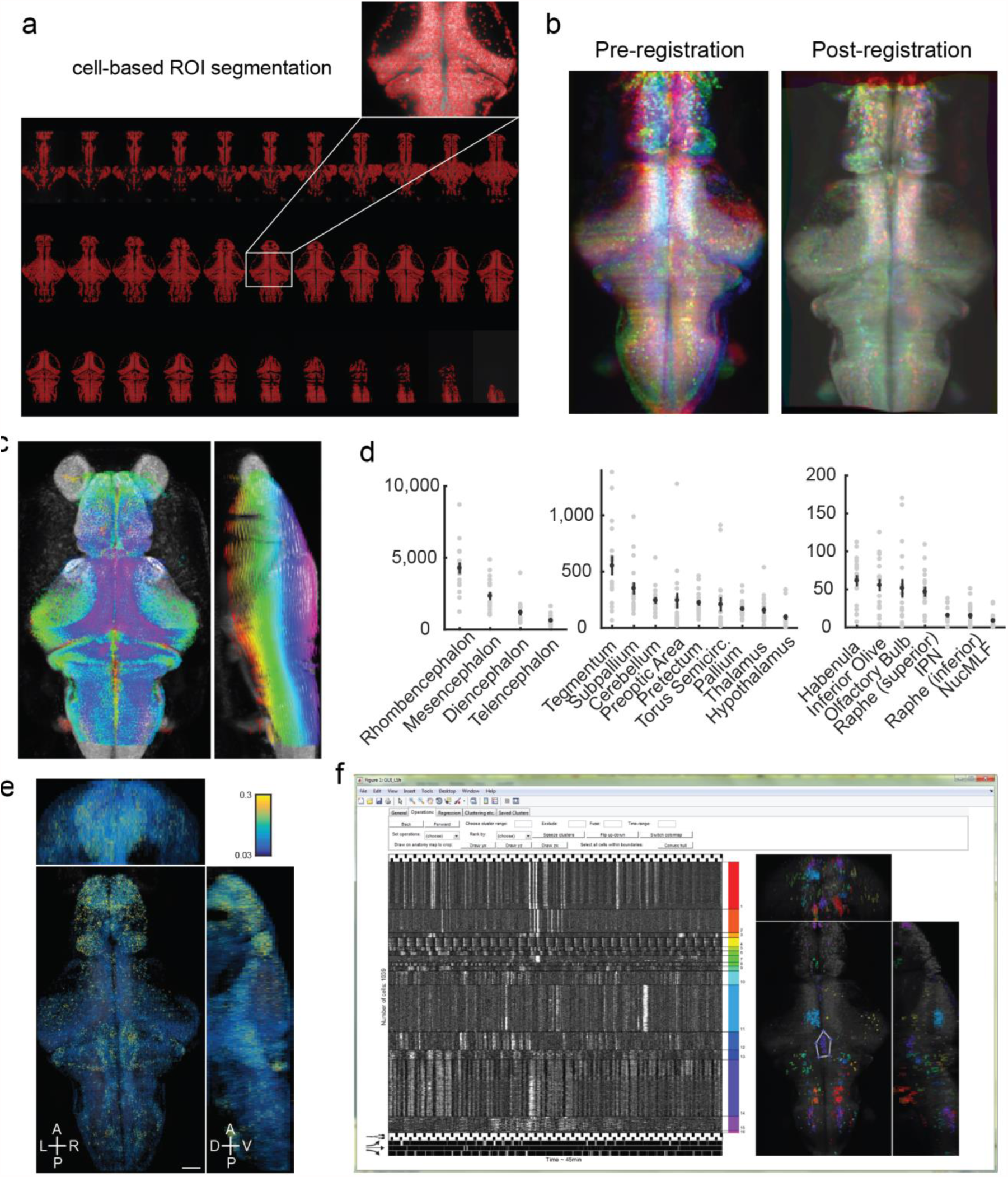
(**a**) Results of automatic cell segmentation based on nucleus-localized calcium indicator. GCaMP6f was expressed panneuronally under the *elavl3* promotor. Inset shows good coverage of segmented ROI’s (red dots) within one imaging plane (total number of segmented ROI’s in this animal: 92,538). (**b**) Overlaid z-projection image of three animals (shown in red/green/blue channels, respectively) before and after registration to an average image stack. (**c**) Registration to the Z-brain reference atlas. Pseudocolors are applied to the z-planes before registration (red through purple: ventral to dorsal). (**d**) Number of cells included in selected anatomical regions according to the Z-brain atlas. Left panel: main brain divisions. Middle and right panel: smaller regions and anatomical features. Grey dots represent data from individual animals (n=18), with mean±SEM shown in black. (**e**) Whole-brain calcium activity averaged per cell over time, shown in pseudo-colors, for one example animal. (**f**) Interface of the custom interactive software that was used to develop the analyses presented in this study. Snapshot shows selected neurons sorted into clusters (indicated by different colors), with their functional activity shown in grayscale in the left panel and their anatomical locations in the right panel.

**Figure S2.**
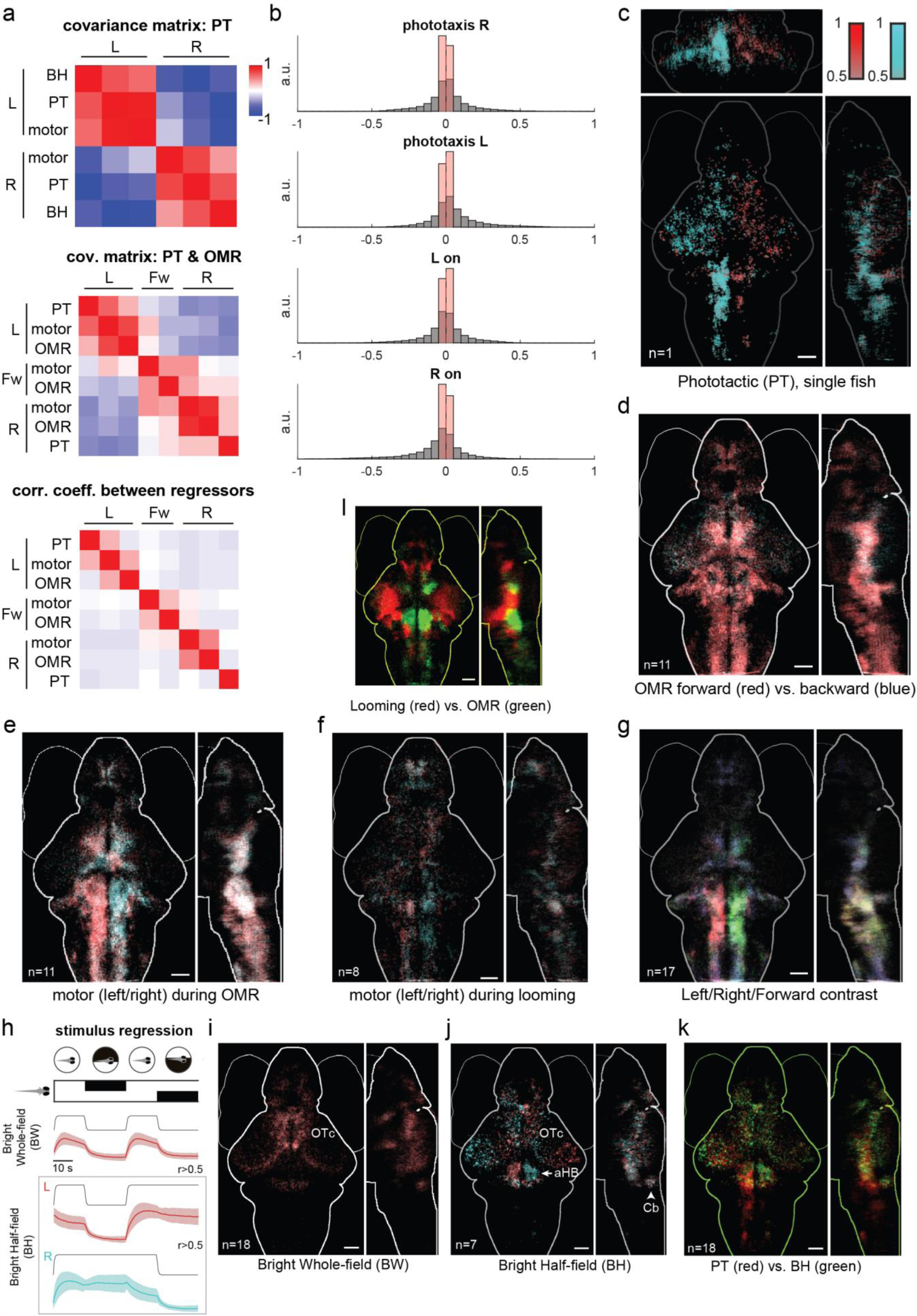
(**a**) Correlation analyses between regressions, for single example fish. Top and middle panel: Covariance matrix or ‘Representational Similarity matrix’ (Kriegeskorte et al., 2008), showing the relationship between whole-brain responses characterized through the various stimulus/motor regressors (see Methods). phT: phototactic stimulus; BH: Bright half-field stimulus; see Fig. S2h. For the middle panel, moving stripes (OMR) were presented in left/right/forward/backward directions, and the fictive recording is parsed into left/right/forward swims regressors. Bottom panel: (direct) correlation coefficients between the regressors (without regressing over cells). This correlation analysis complements the visual comparison of tuning maps that only feature highly correlated neurons. (**b**) Histograms of regression coefficients for all cells from a representative fish, using various regressors. ‘L(R) on’: Bright half-field for the left (right) side being bright. Horizontal axis: Pearson correlation coefficient. (**c**) Single fish example of regression with phototactic regressors. Note that the left side of the hindbrain shows high levels of responses, indicating that leftwards swims are highly correlated with the leftward phototactic stimulus. The finer distinctions between stimulus responses and stimulus-driven motor responses are made in Figure 3. Such asymmetries in behavior (and the corresponding neural correlates) are also typical at the single-animal level (**d**) Motor map during OMR for forward and backward regressors (**e**) Motor map during OMR for left and right regressors. (**f**) Motor map during looming, generated using the behavioral and neural data only during the looming stimulus period. Although the looming stimulus robustly elicits escape responses in freely swimming fish, in tethered preparations there are few behavioral responses. (**g**) Three-way contrast map for left/right/forward swimming. Regression was performed using the 3 fictive swim regressors for left/right/forward (red/green/blue) respectively, and individual cells are colored based on their best regressor. (**h**) Stimulus regressions using phototactic component regressors. Fish were shown a periodic stimulus during imaging that consists of leftwards and rightwards phototactic stimuli separated by a whole-field bright background. The stimulus regressors (black) are constructed by convolving a binary step function with an impulse kernel of GCaMP6. Bright whole-field and bright half-field regressors are shown. See Fig. 2d for phototaxis regression. The colored traces show the dF/F (mean±SD for all ROI’s with r>0.5) for the same example fish as in (a). (**i**) Map for bright whole-field regressor. (**j**) Map for the pair of bright half-field regressors. See Fig. 2e for map for the phototaxis regressors. OTc: optic tectum. aHB: anterior hindbrain. Cb: cerebellum. (**k**) Comparison between phototactic stimulus regression (e, red channel) and bright half-field regression (d, green channel). Yellow indicates overlap between the two color channels. (**l**) Regression map comparing looming (green, same as Fig. 2g) and OMR (red, same as Fig. 2f) regressors.

**Figure S3.**
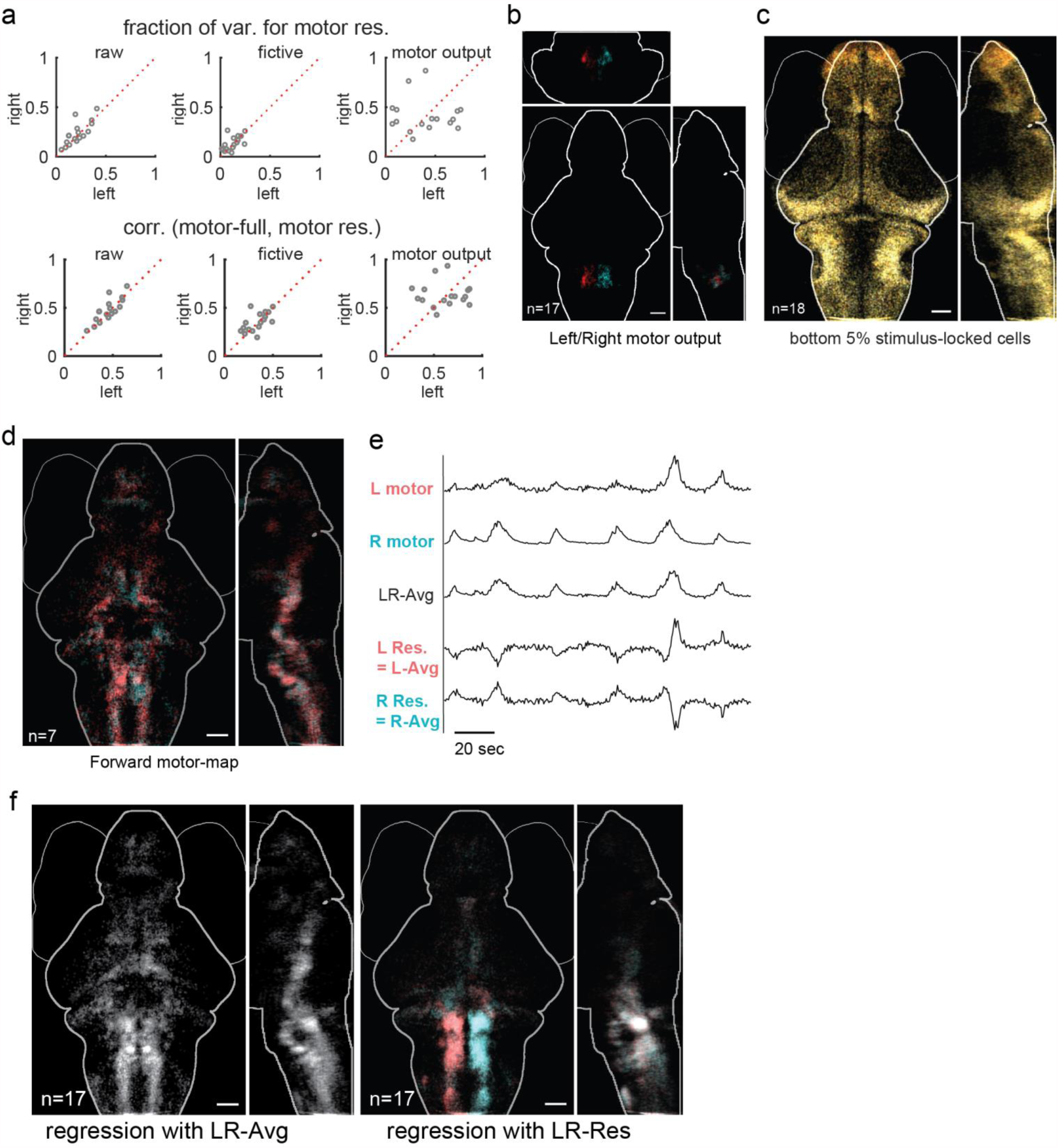
(**a**) Comparison between raw motor recordings, parsed fictive swims, and the ‘motor-seeds’ extracted directed from calcium activity (see Methods). Upper row: fraction of variance explained by the trial-average (tAvr) component of the motor regressor. Lower row: correlation between the full motor regressor and the tAvr component of the motor regressor. The motor outputs showed increased variance explained and correlations, suggesting that they more faithfully represent motor activity patterns. (**b**) Average anatomical map of the cells used to define motor output (ROI’s with highest correlation to the recorded fictive behavior, constrained within Rhombomeres 4 and 5; see Methods). Red: left motor; cyan: right motor. (**c**) Average map of least stimulus-locked (least periodic) cells (compare to Figure 3c). The bottom 5% of cells for each fish were selected, and the average from n=18 fish is shown. (**d**) Forward-swimming map produced by regressing against a pair of forward-swim motor outputs, which were semi-manually extracted from the pair of clusters indicated with red arrows in Fig. 3g. Higher correlation with the left versus right seed is shown in red versus cyan, but the difference between the two seeds is very small. (**e**) Illustration of the left/right motor average and residual. The residual for each side was calculated by subtracting the respective motor activity from the left/right average. **(f**) Related to Fig. 3g; Left: regression map of top 2% of cells ranked by left/right average. Right: regression map of top 2% of cells ranked by left/right residual.

**Figure S4-1.**
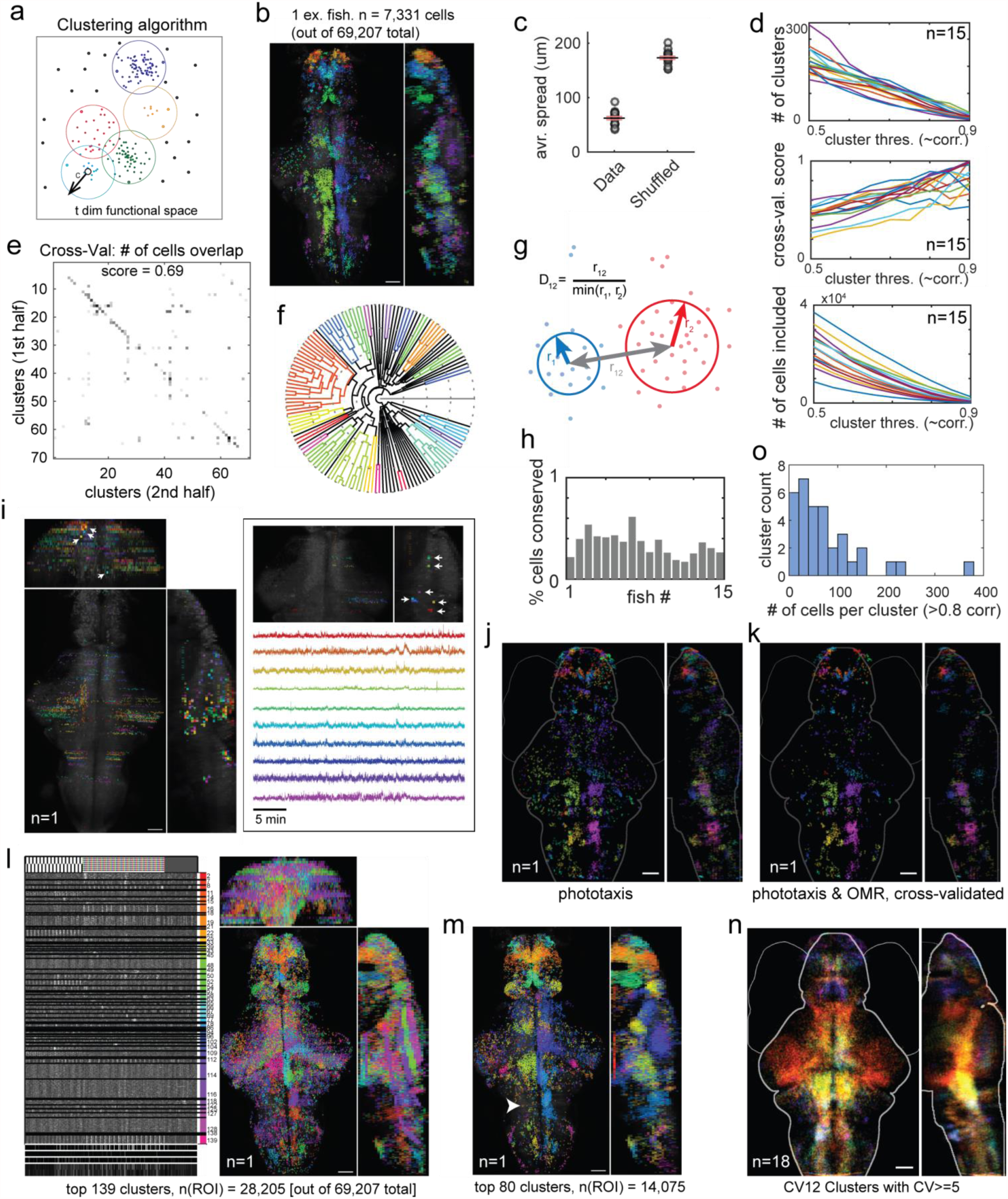
(**a**) Illustration of customized unsupervised clustering algorithm. A density based screen of all cells in functional space was followed by agglomerative clustering with an upper bound on within-cluster dissimilarity (see Methods). (**b**) Results of the automatic clustering algorithm applied to an example fish. The total 139 clusters (6,499 cells) were ordered by hierarchical clustering, and the rainbow colors were assigned based on the resulting leaf order. (**c**) Pearson correlation coefficient matrix between all clusters from (a), calculated with the cluster-average. (**d**) Various clustering statistics as a function of clustering “stringency” threshold (which corresponds to a correlation value), for n = 15 fish (different colored lines). Top: number of clusters. Middle: two-fold cross validation scores, as in (e). Bottom: total number of cells included in all resulting clusters. A threshold of 0.7 was used to obtain results shown in Fig. 4. (**e**) Two-fold cross validation. Clusters produced from the first versus second half of the time-points of the data were matched, and a score was calculated as the fraction of number of cells that were assigned to matched clusters over the total number of cells. This was also visualized as the total “mass” distributed along the diagonal entries. (**f**) Hierarchical clustering diagram of clustering results for an example fish (not the same color code as (b)). (**g**) Illustration of the distance measure used in Fig 4l for assessing whether two clusters were conserved in anatomical space. (**h**) Per fish percentage of clusters (from automatic clustering results) that have anatomically corresponding clusters in at least 6 other fish (out of the 18 fish assessed) as in Fig 4l. (**i**) Clusters from the default clustering (stringency threshold = 0.7) that were identified as artifacts. Inset: for clarity, subset of clusters shown with functional traces. For most of these clusters, the cells within the cluster were aligned along one dimension, e.g. their projections appear as very dense dots (arrows). The dimension corresponds to one of the two laser scanning directions (anterior-posterior, and left-right). A simple script was used to screen out clusters that have very small standard deviation along these physical dimensions. (**j-k**) Cross-validated clusters between two stimuli: phototaxis and OMR (moving gratings). (**j**) All clusters obtained from the phototaxis data from one fish. (**k**) Cross-validation was performed on automatic clustering results for phototaxis versus clustering results for OMR; only cells that were matched in both cross-validation sets were shown, colored according to their original clusters in (j). (**l-m**) Spatial ICA (Hyvärinen and Oja, 2000) as comparison to our clustering method. (**l**) Functional activity and anatomical map of ICA clusters (number of clusters matching that of our clustering method for better comparison). (**m**) A smaller selection of clusters with higher within-cluster correlation shown for clarity. The left half of the hindbrain motor area (arrowhead) is missing. (**n**) Anatomical map of cross-validated clusters (colors ranked as in Fig. 4j), averaged across all fish. Clusters with at least 5 corresponding cells across cross-validation sets were selected. Scale bars, 50 µm. (**o**) Histogram of number of cells within OCM clusters that were very highly correlated to the cluster mean (>0.8 correlation). Results shown for n = 17 fish, with 2 OCM clusters per fish.

**Figure S4-2.**
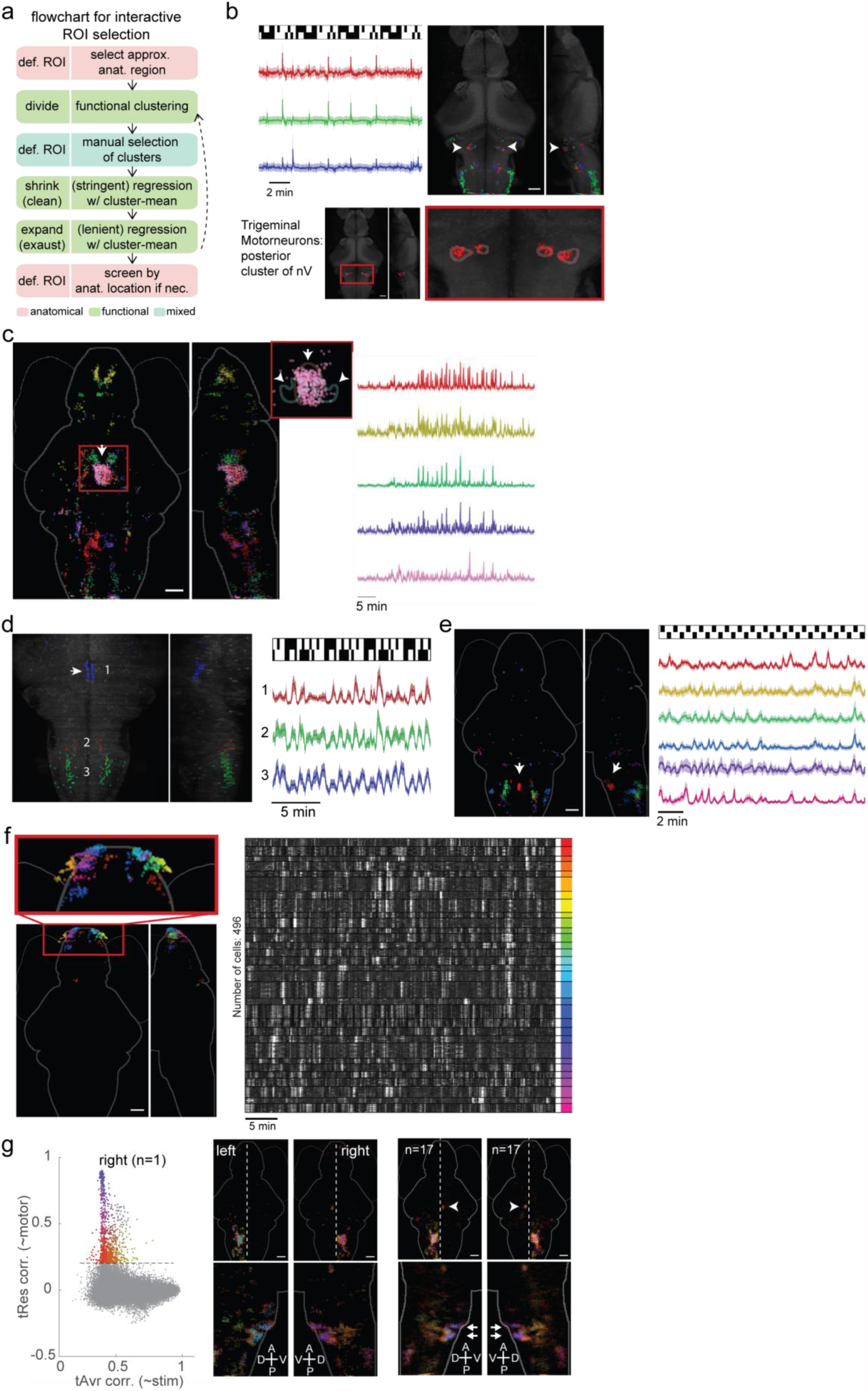
(**a**) Flowchart for interactive ROI selection (using the GUI), showing approximate guidelines. (**b**) Identification of a distinct network related to the trigeminal motor neurons (posterior clusters of nV, arrowheads), characterized by strong, sparse firing (shown in a representative fish). Red/green/blue: network divided into three clusters by k-means that show subtle differences; signal/noise was highest for the green cluster. Inset: anatomical masks for the trigeminal motor neurons from the Z-brain reference atlas - some of these neurons were located within the mask locations. (**c**) Mesencephalic locomotion-related region (pink cluster) and related functional network, shown as a total of 5 k-means clusters (distinct colors). Inset: pink cluster shown with anatomical masks from Z-Brain Atlas. Arrow: (red) mask for “Mesencephalon vglut2 cluster 1”. Arrowheads: pair of (green) masks for Mesencephalon nucMLF (nucleus of the medial longitudinal fascicle). Right: Average functional activity of clusters shown in Fig. S4-2c showing Mesencephalic locomotion-related networks. (**d**) Raphe networks. Left: dorsal raphe nucleus and related networks, shown as 3 k-means clusters. Arrow: dorsal raphe nucleus (as identified by functional clustering). (**e)** Inferior raphe nucleus and related network, shown as 6 k-means clusters. (**f**) Olfactory bulb functional clusters. Left: map of 31 automatically identified clusters shown in different colors (colors assigned according to hierarchical ranking of clusters). Inset: Olfactory bulb region. Clusters corresponding to subnetwork outlined in red in the correlation matrix in Fig. 4i. (**g**) Related to Fig. 4p,q. Left: Two-dimensional sensory-motor plot as in Fig. 4p, showing activity related to right motor output (Fig. 4p is left). Middle: Similar to Fig. 4q, but showing both right and left motor output maps (Fig 4q is the same as the lower left panel). Right: Same analysis as middle, but averaged for n=17 fish. Supplemental Text Accompanying Fig. S4-2.

### Supplemental Text Accompanying Fig. S4-2

#### Jaw and gill movement control

In Fig. S4-2b we show a set of clusters with striking functional characteristics that are found in all animals: extremely sparse and strong activity bouts are observed above a quiet baseline. These clusters were first selected from the whole-brain clustering based on their sparse activity and then extended to include related neurons by regression to their activity. Dividing this larger group into three clusters (k-means, red/green/blue) also revealed subtle differences in functional fingerprints that were consistent across fish, e.g. the caudal population (mainly green) showed a higher signal-to-noise ratio. Some of these neurons were located in the region annotated in the Z-brain atlas as the Trigeminal Motorneurons posterior cluster of nV, while others were found in a characteristically elongated shape in the dorsal-caudal-lateral hindbrain. Previous studies suggest that the trigeminal motor neurons innervate the muscles of the mandibular arch to control jaw movement and are conserved among vertebrates (Higashijima et al., 2000). We hypothesize that these groups of neurons are part of the circuit involved in jaw movement control.

#### Mesencephalonic locomotion-related region

Through our automated clustering, we often observed a prominent cluster in the midbrain (Fig. S4-2c, arrow) that corresponds well to an anatomical mask in the Z-Brain atlas defined from a transgenic line (Mesencephalon vglut2 cluster 1), and is located between the left and right nucMLF (nucleus of the medial longitudinal fascicle). The functional activity of this cluster was highly related to the forward swimming network (see Fig. 3g, S3d, S3f), yet distinct from other clusters within the network (Fig S4-2c, right). Given the role of the nucMLF in controlling forward swimming and that electric stimulation in this area can drive swimming behavior (Severi et al. 2014; electrodes placed near the nucMLF in that experiment), it would be very interesting to further investigate the potential role of this cluster in locomotion generation.

#### Olfactory bulb

When all automatically identified clusters from a single fish were ranked hierarchically, we observed that a relatively large subnetwork of small clusters (adjacent in the hierarchical tree; Fig. 4i, outlined in red) were almost exclusively localized in the olfactory bulb area (Fig. S4-2f left panel), and although their functional activity was not highly correlated between clusters, they shared a similar “texture” in the temporal domain (Fig. S4-2f, right panel). The activity of these clusters was not highly correlated to either the visual input or the motor output. We hypothesize that these clusters represent the functional organization of the olfactory bulb, possibly demonstrating spontaneous activity, given a lack of olfactory stimulation in our experiments (Churchland et al., 2010).

#### Raphe nucleus and the vagus cranial system

When screening for clusters with negative correlation to the motor output, we consistently found an elongated cluster along the midline in the anterior hindbrain that matches the anatomical location of the dorsal raphe nucleus (Fig. S4-2d, left panel). Regression to the raphe activity also consistently revealed two symmetrical groups of neurons in the caudal hindbrain that coincide with the anatomical map of the vagus motor (nX) neurons that control gill movement (Chandrasekhar et al., 1997; Higashijima et al., 2000), suggesting an intriguing functional connection between the two systems that has not to our knowledge been reported before. In future experiments, it would be interesting to verify and interpret this connection between the two systems. These functionally identified networks also often had anatomically well-clustered and symmetrical features, as shown here for the inferior raphe nucleus (Fig. S4-2e).

**Figure S5.**
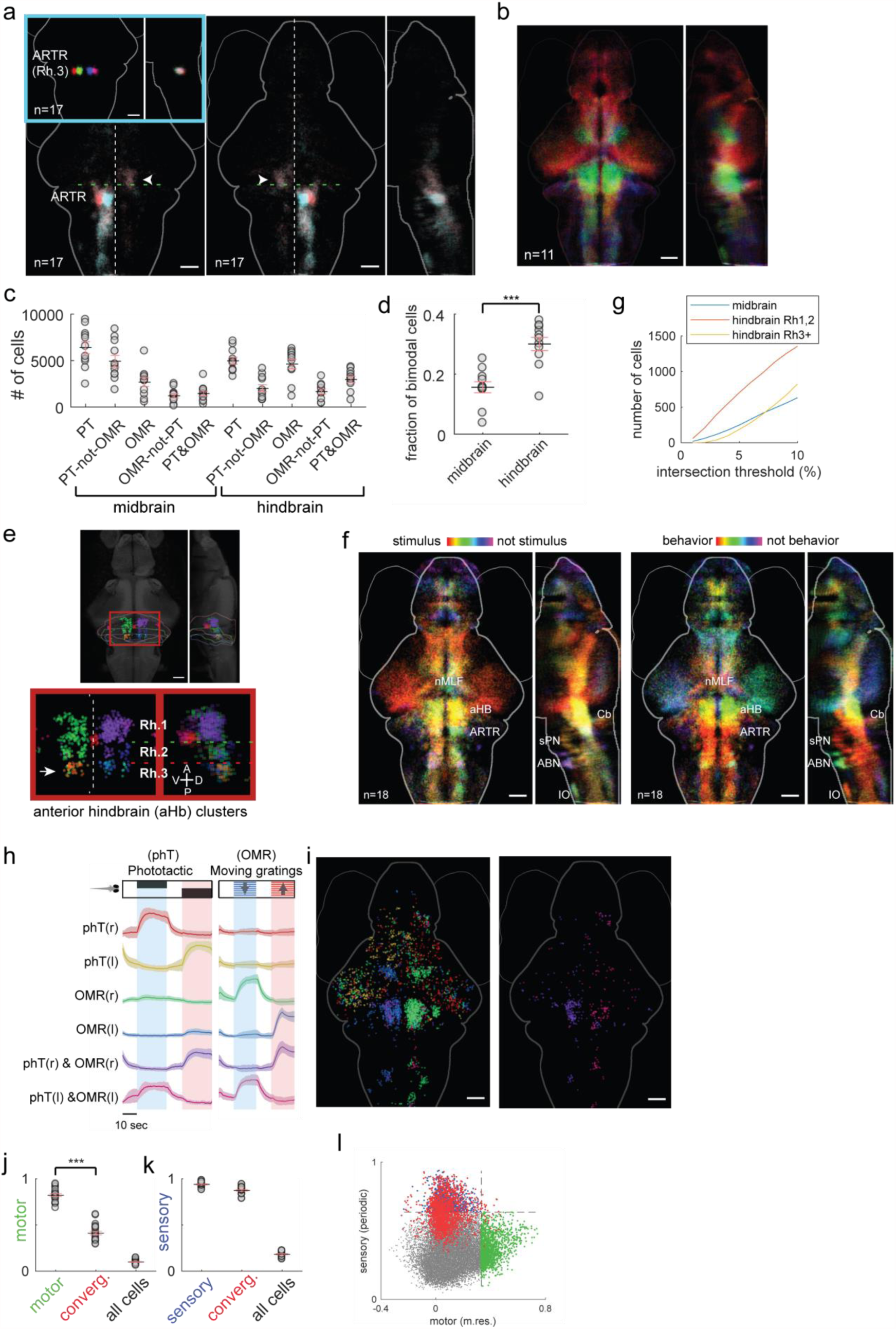
(**a**) Regression maps using the average response of ARTR as regressor (Dunn et al., 2016b). Inset: manually curated ROI’s for ARTR, separating the medial and lateral stripe for each of the left/right side. Arrowhead: regression to the lateral ARTR stripes identifies cells in contralateral Rh.1. (**b**) RGB overlay image of phT only (red), OMR-only (blue) and joint-phT-OMR (blue) cells. Individual cells selected by t-test p<0.001 (leftward-stimulus selectivity contrasted with rightward-stimulus selectivity). (**c**) Quantification of number of unimodal/bimodal cells in the midbrain and hindbrain, respectively, for n=11 fish. Black line: group average. Red lines: standard error of the mean. (**d**) Ratio of bimodal cells over the total number of stimulus selective cells (unimodal plus bimodal). This ratio is significantly higher in the hindbrain (Rh.1 and Rh.2) than in the midbrain (two-tailed t-test, p<0.001). (**e**) Anterior hindbrain clusters as obtained by whole-brain clustering, single fish example. Z-Brain anatomical borders for rhombomeres (Rh) 1-3 are drawn for reference. Arrow: ARTR. (**f**) Left: average map of functional clusters colored by rank as stimulus-locked (i.e. periodic) (red) to not stimulus locked (purple). Right: average map of functional clusters colored by rank for motor res., from most motor related (higher regression coefficients) (red) to least motor related (purple). Similar to single fish map shown in Fig. 4k. Note yellow clusters in the anterior hindbrain that rank relatively highly in both stimulus and behavior. (**g**) Number of convergent cells as a function of threshold (rank %) for (1) midbrain (2) hindbrain Rh1,2 and (3) hindbrain Rh3+. Across the whole range of thresholds, the hindbrain Rh1,2 contains significantly more convergent cells than the midbrain and hindbrain Rh3+ regions. (**h**) Whole-brain regressions were performed to a set of regressors that include phT specific-, OMR specific-, and phT&OMR joint-regressors. For cells that have a correlation coefficient >0.4 to at least one of these regressors, they were classified by their best regressors into 6 groups, color-coded and identified by labels. Average functional activity of these 6 groups of neurons is shown (mean±SD). (**i**) Right panel: anatomical map for single stimulus regressors. Left: anatomical map for convergent regressors. Note that most of these cells are found in Rhombomeres 1&2 of the anterior hindbrain. (**j**) Related to Fig. 5e. Regression coefficient to motor res. for (1) top 5% of cells by motor res. (2) top 5% convergent cells (3) all cells. Corresponds to the x values of the green, red, and gray dots in Fig. 5e, except for all fish and both left and right motor outputs. Note that convergent cells are significantly less motor-related than the most motor-related cells. (**k**) Related to Fig. 5e. (Square root of) variance explained by their periodic component of activity for (1) top 5% of cells by periodicity (2) top 5% convergent cells (3) all cells. Corresponds to the y values of the blue, red, and gray dots in Fig. 5e, except for all fish and both left and right motor outputs. Note that convergent cells are as periodic as the most sensory-related cells. (**l**) Same as Fig. 5e., except for right motor output (Fig. 5e shows data for left motor output). Note that convergent cell activity was less related to right motor output than left. Asymmetric left/right activity patterns and behavioral responses were not uncommon.

